# An innovative optical context to make honeybees crash repeatedly

**DOI:** 10.1101/2021.09.23.461476

**Authors:** Julien R. Serres, Antoine H.P. Morice, Constance Blary, Romain Miot, Gilles Montagne, Franck Ruffier

**Author notes:** **For correspondence:** (JRS). These authors contributed equally to this work (Julien R. Serres, Antoine H.P. Morice, Gilles Montagne, Franck Ruffier). These authors also contributed equally to this work (Constance Blary, Romain Miot).

## Abstract

To investigate altitude control in honeybees, an optical context was designed to make honeybees crash. It has been widely accepted that honeybees rely on the optic flow generated by the ground to control their altitude. However, identifying an optical context capable of uncorrelating forward speed from altitude in honeybees’ flight was the first step towards enhancing the optical context to better understand altitude control in honeybees. This optical context aims to put honeybees in the same flight conditions as an open sky flight above mirror-smooth water. An optical manipulation, based on a pair of opposed horizontal mirrors, was designed to remove any visual information coming from the floor and ceiling. Such an optical manipulation reproduced quantitatively the seminal experiment of Heran & Lindauer (1963), and revealed that honeybees control their altitude by detecting the optic flow with a visual field that extends to approximately 165°.

## Introduction

Flying bees, honeybees or bumblebees, are known to be particularly sensitive to the optic flow pattern generated by the contrasting features of the ground to adjust their altitude by maintaining constant the ventral optic flow during terrain following tasks (***Baird et al., 2006***; ***Portelli et al***., 2010b; ***Srinivasan, 2011***; ***Portelli et al., 2017***; ***Serres and Ruffier, 2017***; ***Lecoeur et al., 2019***), see also (***Franceschini et al., 2007***; ***Portelli et al., 2010a***) for a description of the visuomotor modelling in honeybees. Similarly, honeybees trained to follow the tunnel ceiling while encountering a “dorsal ditch” in the middle of the tunnel configuration (***Portelli et al., 2017***) responded to this new configuration by rising quickly and hugging the new, higher ceiling, by maintaining a similar forward speed, similar distance to the ceiling, and similar dorsal optic flow to those observed during the training step. Conversely, honeybees trained to follow the floor kept on following the floor regardless of the change in the ceiling’s height (***Portelli et al., 2017***).

The present study aims to pursue investigations about the role of dorsal and ventral visual inputs feeding the altitude control system in honeybees by impoverishing the optical context. Inspired by ***Duchon and Warren Jr (2002***) experiments in humans in which they designed an optical manipulation made with a pair of infinite walls in order to optically remove the floor, then by strongly impoverishing visual information coming from the floor, we designed a novel optical context to make honeybees crash irremediably into the floor. Such an optical manipulation in which the floor appeared to be removed will allow us to quantitatively reproduce and extend the seminal experiment of ***Heran and Lindauer (1963***). Sixty years ago, these authors trained honeybees to fly above a 247m-long water surface. When the water surface was rippled or when a floating bridge provided a visual contrast, honeybees were able to cross the lake. However, honeybees crossing mirror-smooth water during foraging trips flew lower and lower until they crashed and drowned (***Heran and Lindauer, 1963***). To replicate and extend experimentally such a behaviour observed outdoor above a water surface in a flight tunnel, we used a pair of mirrors placed on the floor and on the ceiling.

## Methods and Materials

### Flight tunnel

The flight tunnel (Fig. 2) is a rectangular shape (220cm long, 71cm high and 25cm wide), in which a unique pattern is printed on every tunnel’s wall and reproduced with red gelatin filter’s stripes (Lee Filters HT019) on the right wall. The pattern on the four surfaces of the tunnel consists of red and white stripes oriented perpendicularly to the insect’s flight path. The entrance is circular (5 cm in diameter) and located at 9-14 cm from the floor. This video can be watched to better understand the organization of our experimental set-up: https://youtu.be/KH9z8eqOBbU.

### Pattern

These red stripes of two different widths (1 cm and 3 cm) form a simple 10 cm-wide pattern regularly repeated, visible in Fig. 2. The angular subtends of the stripes ranged from 5.7° to 53° (1–10 cm-wide pattern viewed from a distance of 10 cm, respectively) and from 0.5° to 5.3° (1–10 cm-wide pattern viewed from 1 m, respectively). As honeybees do not possess red-sensitive photoreceptors (***Srinivasan, 2011***), they perceive red stripes as grey ones. Between the red and white stripes, the Michelson contrast is 0.47 on the planks and 0.25 on the insect netting. Contrast was measured using a photodiode equipped with a green band-pass filter (Kodak Wratten n°61), the transmission spectrum of which closely matched the spectral sensitivity of the honeybee’s green receptors.

### Experimental procedure

During the experimental period, a small set of four groups comprising from 14 to 27 freely flying honeybees (*Apis mellifera*) were color-marked and trained daily to enter and fly alone along the outdoor tunnel to collect a sugar solution reward (strength by mass: 25% sucrose) at the opposite end. Their flight path toward the reward was recorded with a digital camera from the insect-netting side, strictly in keeping with the chronology of the procedure depicted in Fig. 1 using a distinct group of honeybees for each of the four experimental conditions (A, B, C, and D).

**Figure 1.**
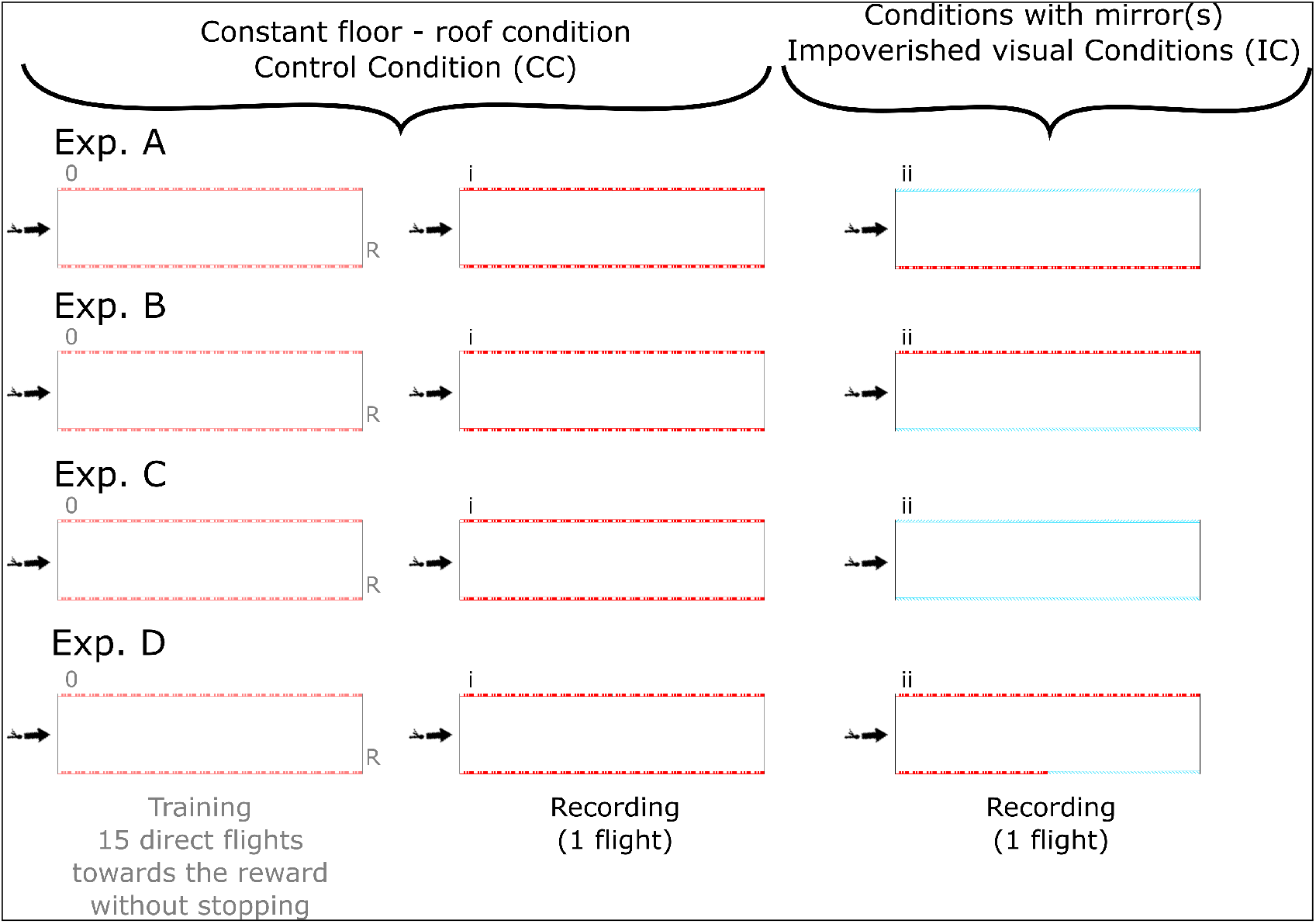
Chronology of the procedure used in the four experimental conditions (A,B,C, and D). The same individual bees were trained and tested and their trajectories were recorded in line with the strict chronology of the steps (0, i, and ii). **Step 0 (the training step - 15 direct flights):** the honeybees were first trained by completing about 15 flights travelling along the tunnel with a uniform height of 71cm (called “Constant floor-roof condition”) to collect nectar in a reward box. Both the entrance and the reward were placed near the floor. Honeybees were rewarded at the end of each flight; the entrance to the reward box was closed during the flights so that no visual cues were available about the position of the reward. **Step i (Control Condition CC, video-recording - 1 flight):** immediately after the training step, individual bees’ trajectories were recorded in this same tunnel with a uniform height of 71cm (called “Constant floor-roof condition”); the entrance to the reward box was closed during the flights so that no visual cues were available about the position of the reward. **Step ii (Impoverished visual Conditions IC, video-recording - 1 flight):** immediately after recording the bees’ trajectories under “Constant floor-roof condition”, one mirror (Experimental conditions A & B), two mirrors (Experimental condition C), or the half low mirror (Experimental condition D) were uncovered, and the bees’ trajectories were recorded in the presence of an optical manipulation (either virtually doubling the tunnel height when there is one mirror uncovered, or giving virtually an infinite tunnel height when there are two mirrors); the entrance to the reward box was again closed during these flights so that no visual cues were available about the position of the reward.

**Figure 2.**
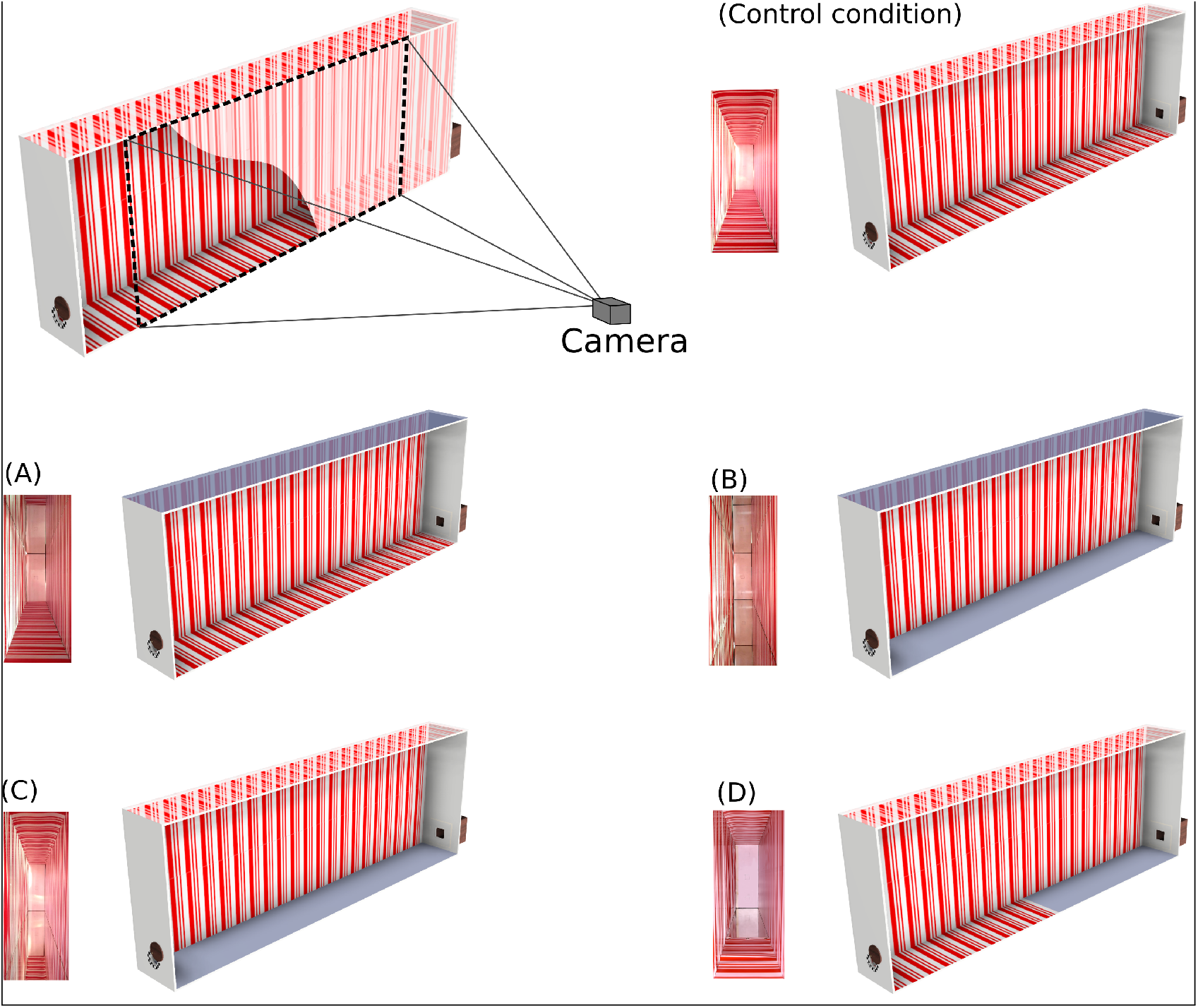
The flight tunnel was a rectangular shape (220cm long,71cm high and 25cm wide. In the control condition (CC), the four surfaces of the tunnel were textured with red and white stripes oriented perpendicularly to the bees’ flight paths. One side of the tunnel consisted of insect netting lined with stripes. The camera was placed sideways, 2.3 m from the insect netting. The field of view (130 cm in width, 71 cm in height) covered the whole height of the tunnel, from abscissa x = 0 cm to abscissa x = 130cm in experiments (A-C), and from abscissa x = 35cm to abscissa x = 165cm in experiment (D). In experiment (A), only the top mirror was uncovered. In experiment (B), both mirrors were uncovered. In experiment (C), only the bottom mirror was uncovered. In experiment (D), only the bottom mirror was half uncovered.

In the first session (called “Constant floor-roof condition” - Control Condition CC), honeybees were trained during 16 trials to follow the ground (Fig. 1A(0+i)-D(0+i)). In a second session (called “Conditions with mirror(s)” - Impoverished visual Conditions IC), one mirror, two mirrors, or the half low mirror were uncovered (Fig. 1A(ii)-D(ii)), appearing to double the tunnel’s height when there was one (or half) mirror uncovered (above in Exp. A; below in Exp. B & D), or giving the appearance of an infinite tunnel height when there were two mirrors uncovered in Exp. C.

### Video recordings and flight path analysis

The honeybees’ trajectories were filmed at a rate of 100 frames per second with a high-resolution black-and-white CMOS camera (Teledyne Dalsa Genie HM640). The video recording was manually triggered after the experimenter opened the tunnel entrance to the honeybees, who were sent through one by one. A red filter (Lee Filters HT019) is set in front of the camera monitoring honeybee’s traces. This process removes the red stripes on the trajectory records and optimizes the contrast between the honeybee and the background. The camera was placed sideways, 2.3m from the insect netting. The field of view (130cm in width, 71cm in height) covered the whole height of the tunnel, from abscissa x = 0cm to abscissa x = 130cm in experiments A-C, and from abscissa x = 35cm to abscissa x = 165cm in experiment D. Image sequences were calibrated, corrected, processed and analysed using a custom-made Matlab program available online (see: https://github.com/rm1720/bees-applications/wiki). This program automatically determined the hon-eybees’ flight height *z* in each frame as a function of the abscissa *x* along the tunnel axis so that the honeybee’s trajectory in the vertical plane could be plotted. Only trajectories until the first crash were plotted and analysed.

### Statistical analysis

All the data recorded were included in the statistical analysis without removing any outliers. Statistical data analyses were performed with the Matlab R2018a software program. Not all dataset exhibit the same number of measurements as a function of the abscissa because the bees’ crash creates an incomplete trajectory. As a consequence, median and Median Absolute Deviation (Median ± MAD) values were computed for all datasets by binning each bee’s trajectory with an 8cm window in abscissa (see Table 1). Mann–Whitney U tests are used to compare altitude binning distribution by pairs : control conditions (CC) versus impoverished visual conditions (IC) in each experiment (A, B, C, and D) (see Table 1).

**Table 1.**
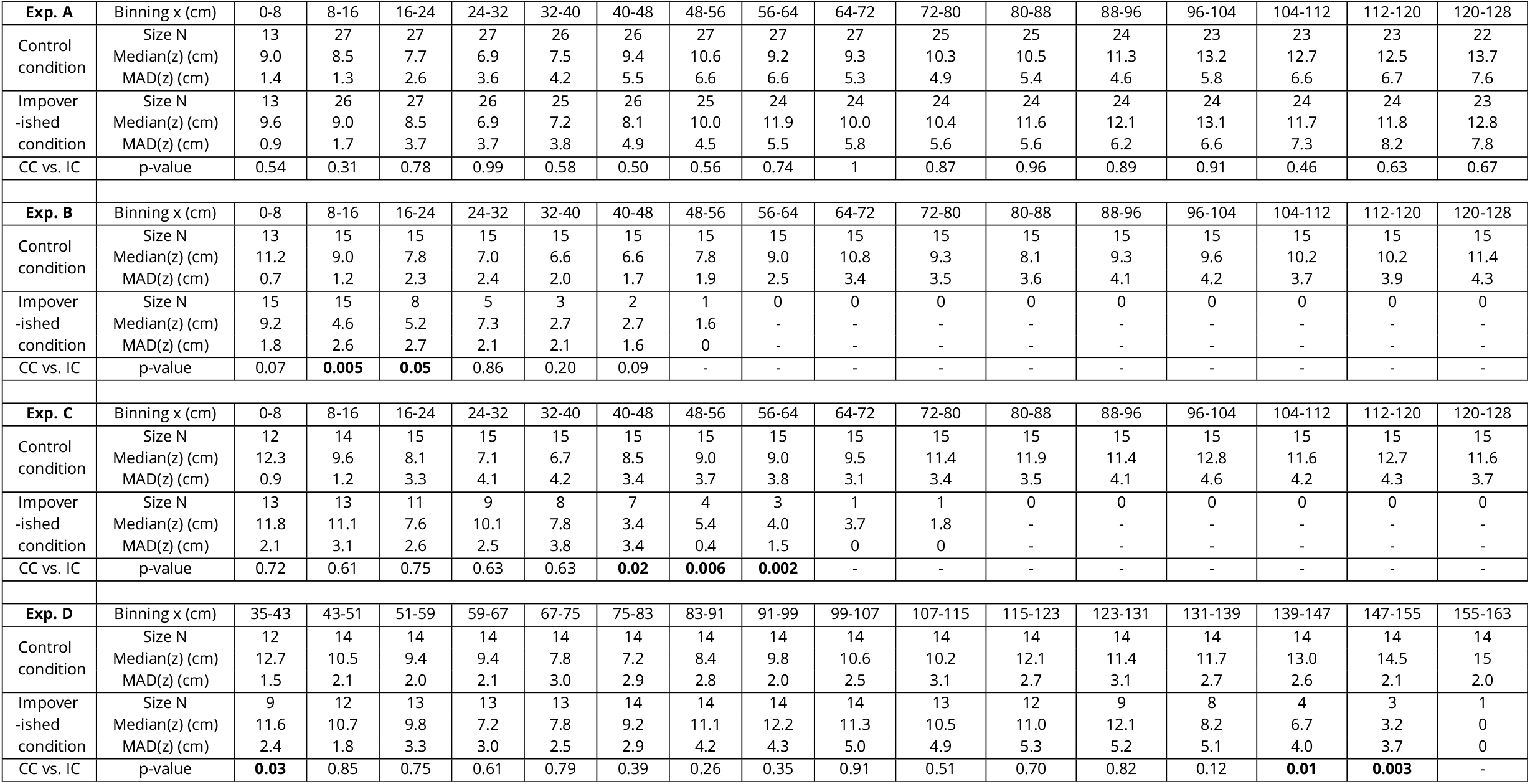
Comparison of the altitude distributions in each experiment (A, B, C, and D) with a binning of 8 cm along the abscissa. For each experiment and both conditions (control condition CC and impoverished condition IC), the size of the sample in the binning (N), the median, and Median Absolute Deviation (MAD) are given. Mann–Whitney U tests are used to compare altitude binning distribution by pairs : control conditions (CC) versus impoverished visual conditions (IC) in each experiment (A, B, C, and D). Significant probabilities (CC vs. IC) are in bold.

## Results

### Honeybees following the floor do not rely on dorsal visual information

In experiment A (Fig. 3A), we tested the effect of a visual impoverishment in the dorsal part of the honeybees’ visual field. The chronology of the procedure from step 0 to step ii is described in Fig. 1. A group of 27 honeybees was trained in the control condition (CC) (Fig. 3Ai-Aii). The 1st flight “in mirror on the ceiling” condition, in which the top mirror was uncovered, was recorded in impoverished visual condition (IC) (See Experimental procedure). The presence of the mirror on the ceiling appearing to double the tunnel’s height (142cm) upwards (Fig. 3Aiii-Aiv). We observed no significant change in flight behaviour in honeybees (Exp. A in Table 1).

**Figure 3.**
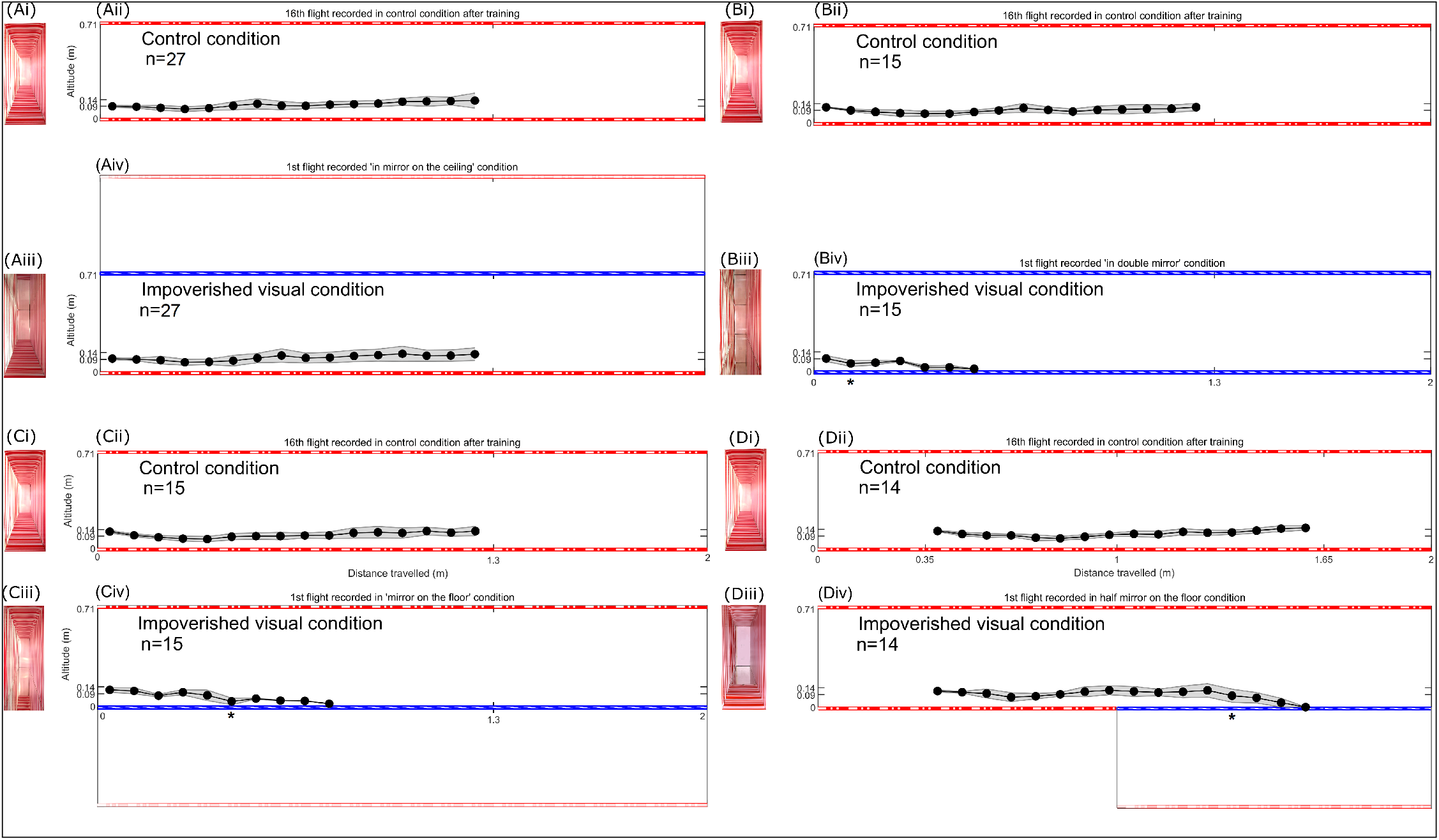
This figure depicts the trajectories (Median ± MAD) followed by honeybees in the vertical plane (*x,z*) in the four experimental conditions (A,B,C, and D). In each panel, a comparison can be made between the trajectories produced prior to (Aii, Bii, Cii and Dii) and after (Aiv, Biv, Civ and Div) experimental manipulations. These manipulations consisted of adding a mirror either on the ceiling (Aiii), on both the floor and the ceiling (Biii), on the floor (Ciii) or on half the floor (Diii). Results show that while manipulating dorsal optic flow doesn’t affect the honeybees’ trajectories, manipulating ventral optic flow, whether it is a simple manipulation or a suppression, has strong consequences on the trajectories, leading to systematic crashes.

### Without any ventral and dorsal visual information, honeybees crash irremediably

In experiment B (Fig. 3B), we tested the effect of a visual impoverishment in both the dorsal and ventral parts of the honeybees’ visual field. The chronology of the procedure is described in Fig. 1. A group of 15 honeybees was trained in the control condition (CC) (Fig. 3Bi-Bii). The 1st flight, “in double mirror” condition in which both mirrors were uncovered, was recorded in impoverished visual condition (IC) (See Experimental procedure). The presence of mirrors on the top and on the bottom created an optical manipulation in which a pair of infinite walls appeared. As a result, no visual information from either the floor or ceiling was available (Fig. 3Biii-Biv). We observed significant changes in flight behaviour in honeybees (Exp. B in Table 1) from x = 8cm until all the honeybees crashed into the floor.

### Dorsal visual information does not aid in flying further before crashing

In experiment C (Fig. 3C), we tested the effect of a visual impoverishment in the ventral part of the honeybees’ visual field. The chronology of the procedure is described in Fig. 1. A group of 15 honeybees was trained in the control condition (CC) (Fig. 3Ci-Cii). The 1st flight “in mirror on the floor” condition in which the bottom mirror was uncovered, was recorded in impoverished visual condition (IC) (See Experimental procedure). The presence of the mirror on the floor appearing to double the tunnel’s height (142cm) downwards (Fig. 3Ciii-Civ) making a kind of “ventral ditch” of 71cm in depth. We observed significant changes in flight behaviour in honeybees (Exp. C in Table 1) from x = 40cm until all the honeybees crashed into the floor. Honeybees may be visually attracted by the virtual floor 71cm below, but then they crash into the mirror on the floor.

### Adding a sheet of texture does not help honeybees to fly further above the mirror

In experiment D (Fig. 3D), we tested the effect of a visual enhancement in the ventral part of the honeybees’ visual field by adding on the first half of the bottom mirror a sheet of texture. The chronology of the procedure is described in Fig. 1. A group of 14 honeybees was trained in the control condition (CC) (Fig. 3Ci-Cii). The 1st flight “in half mirror on the floor” condition in which the bottom mirror was half uncovered, doubling virtually the tunnel height (142cm) downwards (Fig. 3Diii-Div) making a kind of “ventral ditch” of 71cm in depth as in experiment C (Fig. 3C). We observed significant changes in flight behaviour in honeybees (Exp. C in Table 1) from x = 139cm until all honeybees crashed into the floor. This additional sheet of texture does not help honeybees to fly further above the bottom mirror, the flight distance above the mirror (39cm), where we observed a significant difference in height, was similar to the one observed in experiment C (Table 1). The average height of flight in experiment C was 10.2cm (position marked by a * symbol in Fig. 3C) before losing altitude (Table 1) which allowed computation of the angular position of the texture - mirror at ∼ 165° ventrally 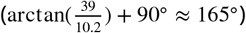.

## Discussion and conclusion

The aim of the present study was to extend ***Heran and Lindauer (1963***) work by precisely manipulating parts of the optic flow available. The use of mirrors that could cover all or part of the upper and lower visual field, allowed us to examine the influence of the deprivation of several parts of optic flow on honeybees’ trajectories. Developing such an experimental set-up will subsequently permit enhancement of the optical context to thoroughly investigate altitude control in honeybees.

In experiment A (Fig. 3Aiv), honeybees appear to follow the floor despite this virtual “dorsal ditch” because they may have learnt to follow the floor and to rely on the ventral visual information in order to regulate their flight and to forage, in harmony with ***Portelli et al. (2017***) results.

Conversely, each experimental manipulation that affects the ventral part of the optic flow, whether it is a total deprivation of the ventral optic flow in experiment C (Fig. 3Biv)) or a change in the resulting the ventral part of the optic flow (Figs. 3Civ,Div), gave rise to crashing into the mirror. Interestingly our double mirror condition allowed us to replicate the untextured experimental condition used by ***Heran and Lindauer (1963***) in a naturalistic environment. Our results are in perfect agreement with theirs insofar as the honeybees crash consistently in the absence of ventral optic flow.

In experiments C and D, the use of a mirror on the floor does not cause the suppression of the ventral part of optic flow, but the ceiling textures mirrored in the floor create a virtual “ventral gap” suppressing the ventral part of optic flow. The reduction in honeybees’ altitude until the crash could result from a change in altitude intended to restore the optically specified altitude as experienced in the 15 control trials, which is in harmony with both ***Portelli et al. (2010b***) and ***Portelli et al. (2017***) results. Taken together, all our results indicate unequivocally that the crashes observed in the study of ***Heran and Lindauer (1963***) reflect the propensity of honeybees to reduce their altitude, in order to restore the ventral optic flow experienced by honeybees in equivalent tasks.

Results in experiment D (Fig. 3Div) reveal that honeybees regulate the ventral optic flow with a visual field that extends to approximately 165° ventrally. By picking up information over a wide visual field (***Lecoeur et al., 2019***), honeybees detect and respond to changes in the environment even when lacking texture on the ground.

## Supporting information

Data set of honeybees flights

## Acknowledgments

The authors would like to thank Marc Boyron and Julien Diperi for their technical assistance with designing the flight tunnel, Patrick Sainton for his video montage showing the experimental set-up, and they wish to thank David Wood (English at your Service, http://www.eays.eu/) for revising the English of the manuscript.

## Additionnal information

### Author contributions

JRS, AHPM, GM, and FR designed the experiments. CB performed the experiments. RM developed the tracking app with Matlab software. JRS, AHPM, CB, GM, and FR analysed the data. JRS wrote the first draft of the paper; all authors prepared and revised the manuscript. JRS coordinated and supervised this research work.

### Ethics

Animal experimentation: study involved experiments on honeybees and were conducted according to ethical guidelines.

## References

Baird E, Srinivasan MV, Zhang S, Lamont R, Cowling A. Visual control of flight speed and height in the honeybee. In: International Conference on Simulation of Adaptive Behavior Springer; 2006. p. 40–51.

Duchon AP, Warren Jr WH. A visual equalization strategy for locomotor control: of honeybees, robots, and humans. Psychological Science. 2002; 13(3):272–278.

Franceschini N, Ruffier F, Serres J. A bio-inspired flying robot sheds light on insect piloting abilities. Current Biology. 2007; 17(4):329–335.

Heran H, Lindauer M. Windkompensation und seitenwindkorrektur der bienen beim flug über wasser. Zeitschrift für vergleichende Physiologie. 1963; 47(1):39–55.

Lecoeur J, Dacke M, Floreano D, Baird E. The role of optic flow pooling in insect flight control in cluttered environments. Scientific reports. 2019; 9(1):1–13.

Portelli G, Serres J, Ruffier F, Franceschini N. Modelling honeybee visual guidance in a 3-D environment. Journal of Physiology - Paris. 2010; 104(1-2):27–39.

Portelli G, Ruffier F, Franceschini N. Honeybees change their height to restore their optic flow. Journal of Comparative Physiology A. 2010; 196(4):307–313.

Portelli G, Serres JR, Ruffier F. Altitude control in honeybees: joint vision-based learning and guidance. Scientific reports. 2017; 7(1):1–10.

Serres JR, Ruffier F. Optic flow-based collision-free strategies: From insects to robots. Arthropod structure & development. 2017; 46(5):703–717.

Srinivasan MV. Honeybees as a model for the study of visually guided flight, navigation, and biologically inspired robotics. Physiological reviews. 2011; 91(2):413–460.

