## Supplementary material for "An innovative optical context to make honeybees crash repeatedly": Data set of honeybees flights: binning.m

```
function [mean_Z]=binning(pas,X,Z)
%pas = 0.1;
taille = 1.3/pas;
for k=0:(taille-1)
  index=find(k*pas<X & X<(k+1)*pas);
  if length(index)~=0
  mean_Z(k+1) = mean(Z(index));
  end
  if length(index)==0
      mean_Z(k+1)=NaN;
  end
  clear index
  %pause
end
```
