## Supplementary material for "An innovative optical context to make honeybees crash repeatedly": Data set of honeybees flights: rgb.m

```
% RGB  Rgb triple for given CSS color name
%
%   RGB = RGB('COLORNAME') returns the red-green-blue triple corresponding
%     to the color named COLORNAME by the CSS3 proposed standard [1], which
%     contains 139 different colors (an rgb triple is a 1x3 vector of
%     numbers between 0 and 1). COLORNAME is case insensitive, and for gray
%     colors both spellings (gray and grey) are allowed.
%
%   RGB CHART creates a figure window showing all the available colors with
%     their names.
%
%   EXAMPLES
%     c = rgb('DarkRed')               gives c = [0.5430 0 0]
%     c = rgb('Green')                 gives c = [0 0.5 0]
%     plot(x,y,'color',rgb('orange'))  plots an orange line through x and y
%     rgb chart                        shows all the colors
%
%   BACKGROUND
%     The color names of [1] have already been ratified in [2], and
%     according to [3] they are accepted by almost all web browsers and are
%     used in Microsoft's .net framework. All but four colors agree with
%     the X11 colornames, as detailed in [4]. Of these the most important
%     clash is green, defined as [0 0.5 0] by CSS and [0 1 0] by X11. The
%     definition of green in Matlab matches the X11 definition and gives a
%     very light green, called lime by CSS (many users of Matlab have
%     discovered this when trying to color graphs with 'g-'). Note that
%     cyan and aqua are synonyms as well as magenta and fuchsia.
%
%   ABOUT RGB
%     This program is public domain and may be distributed freely.
%     Author: Kristj�n J�nasson, Dept. of Computer Science, University of
%     Iceland. June 2009.
%
%   REFERENCES
%     [1] "CSS Color module level 3", W3C (World Wide Web Consortium)
%         working draft 21 July 2008, http://www.w3.org/TR/css3-color
%
%     [2] "Scalable Vector Graphics (SVG) 1.1 specification", W3C
%         recommendation 14 January 2003, edited in place 30 April 2009,
%         http://www.w3.org/TR/SVG
%
%     [3] "Web colors", http://en.wikipedia.org/wiki/Web_colors
%
%     [4] "X11 color names" http://en.wikipedia.org/wiki/X11_color_names

function rgb = rgb(s)
  persistent num name
  if isempty(num) % First time rgb is called
    [num,name] = getcolors();
    name = lower(name);
    num = reshape(hex2dec(num), [], 3);
    % Divide most numbers by 256 for "aesthetic" reasons (green=[0 0.5 0])
    I = num < 240;  % (interpolate F0--FF linearly from 240/256 to 1.0)
    num(I) = num(I)/256;
    num(~I) = ((num(~I) - 240)/15 + 15)/16; + 240;
  end
  if strcmpi(s,'chart')
    showcolors()
  else
    k = find(strcmpi(s, name));
    if isempty(k)
      error(['Unknown color: ' s]);
    else
      rgb = num(k(1), :);
    end
  end
end

function showcolors()
  [num,name] = getcolors();
  grp = {'White', 'Gray', 'Red', 'Pink', 'Orange', 'Yellow', 'Brown'...
    , 'Green', 'Blue', 'Purple', 'Grey'};
  J = [1,3,6,8,9,10,11];
  fl = lower(grp);
  nl = lower(name);
  for i=1:length(grp)
    n(i) = strmatch(fl{i}, nl, 'exact'); 
  end
  clf
  p = get(0,'screensize');
  wh = 0.6*p(3:4);
  xy0 = p(1:2)+0.5*p(3:4) - wh/2;
  set(gcf,'position', [xy0 wh]);
  axes('position', [0 0 1 1], 'visible', 'off');
  hold on
  x = 0;
  N = 0;
  for i=1:length(J)-1
    N = max(N, n(J(i+1)) - n(J(i)) + (J(i+1) - J(i))*1.3); 
  end
  h = 1/N;
  w = 1/(length(J)-1);
  d = w/30;
  for col = 1:length(J)-1;
    y = 1 - h;
    for i=J(col):J(col+1)-1
      t = text(x+w/2, y+h/10 , [grp{i} ' colors']);
      set(t, 'fontw', 'bold', 'vert','bot', 'horiz','cent', 'fontsize',10);
      y = y - h;
      for k = n(i):n(i+1)-1
        c = rgb(name{k});
        bright = (c(1)+2*c(2)+c(3))/4;
        if bright < 0.5, txtcolor = 'w'; else txtcolor = 'k'; end
        rectangle('position',[x+d,y,w-2*d,h],'facecolor',c);
        t = text(x+w/2, y+h/2, name{k}, 'color', txtcolor);
        set(t, 'vert', 'mid', 'horiz', 'cent', 'fontsize', 9);
        y = y - h;
      end
      y = y - 0.3*h;
    end
    x = x + w;
  end
end

function [hex,name] = getcolors()
  css = {
    %White colors
    'FF','FF','FF', 'White'
    'FF','FA','FA', 'Snow'
    'F0','FF','F0', 'Honeydew'
    'F5','FF','FA', 'MintCream'
    'F0','FF','FF', 'Azure'
    'F0','F8','FF', 'AliceBlue'
    'F8','F8','FF', 'GhostWhite'
    'F5','F5','F5', 'WhiteSmoke'
    'FF','F5','EE', 'Seashell'
    'F5','F5','DC', 'Beige'
    'FD','F5','E6', 'OldLace'
    'FF','FA','F0', 'FloralWhite'
    'FF','FF','F0', 'Ivory'
    'FA','EB','D7', 'AntiqueWhite'
    'FA','F0','E6', 'Linen'
    'FF','F0','F5', 'LavenderBlush'
    'FF','E4','E1', 'MistyRose'
    %Grey colors'
    '80','80','80', 'Gray'
    'DC','DC','DC', 'Gainsboro'
    'D3','D3','D3', 'LightGray'
    'C0','C0','C0', 'Silver'
    'A9','A9','A9', 'DarkGray'
    '69','69','69', 'DimGray'
    '77','88','99', 'LightSlateGray'
    '70','80','90', 'SlateGray'
    '2F','4F','4F', 'DarkSlateGray'
    '00','00','00', 'Black'
    %Red colors
    'FF','00','00', 'Red'
    'FF','A0','7A', 'LightSalmon'
    'FA','80','72', 'Salmon'
    'E9','96','7A', 'DarkSalmon'
    'F0','80','80', 'LightCoral'
    'CD','5C','5C', 'IndianRed'
    'DC','14','3C', 'Crimson'
    'B2','22','22', 'FireBrick'
    '8B','00','00', 'DarkRed'
    %Pink colors
    'FF','C0','CB', 'Pink'
    'FF','B6','C1', 'LightPink'
    'FF','69','B4', 'HotPink'
    'FF','14','93', 'DeepPink'
    'DB','70','93', 'PaleVioletRed'
    'C7','15','85', 'MediumVioletRed'
    %Orange colors
    'FF','A5','00', 'Orange'
    'FF','8C','00', 'DarkOrange'
    'FF','7F','50', 'Coral'
    'FF','63','47', 'Tomato'
    'FF','45','00', 'OrangeRed'
    %Yellow colors
    'FF','FF','00', 'Yellow'
    'FF','FF','E0', 'LightYellow'
    'FF','FA','CD', 'LemonChiffon'
    'FA','FA','D2', 'LightGoldenrodYellow'
    'FF','EF','D5', 'PapayaWhip'
    'FF','E4','B5', 'Moccasin'
    'FF','DA','B9', 'PeachPuff'
    'EE','E8','AA', 'PaleGoldenrod'
    'F0','E6','8C', 'Khaki'
    'BD','B7','6B', 'DarkKhaki'
    'FF','D7','00', 'Gold'
    %Brown colors
    'A5','2A','2A', 'Brown'
    'FF','F8','DC', 'Cornsilk'
    'FF','EB','CD', 'BlanchedAlmond'
    'FF','E4','C4', 'Bisque'
    'FF','DE','AD', 'NavajoWhite'
    'F5','DE','B3', 'Wheat'
    'DE','B8','87', 'BurlyWood'
    'D2','B4','8C', 'Tan'
    'BC','8F','8F', 'RosyBrown'
    'F4','A4','60', 'SandyBrown'
    'DA','A5','20', 'Goldenrod'
    'B8','86','0B', 'DarkGoldenrod'
    'CD','85','3F', 'Peru'
    'D2','69','1E', 'Chocolate'
    '8B','45','13', 'SaddleBrown'
    'A0','52','2D', 'Sienna'
    '80','00','00', 'Maroon'
    %Green colors
    '00','80','00', 'Green'
    '98','FB','98', 'PaleGreen'
    '90','EE','90', 'LightGreen'
    '9A','CD','32', 'YellowGreen'
    'AD','FF','2F', 'GreenYellow'
    '7F','FF','00', 'Chartreuse'
    '7C','FC','00', 'LawnGreen'
    '00','FF','00', 'Lime'
    '32','CD','32', 'LimeGreen'
    '00','FA','9A', 'MediumSpringGreen'
    '00','FF','7F', 'SpringGreen'
    '66','CD','AA', 'MediumAquamarine'
    '7F','FF','D4', 'Aquamarine'
    '20','B2','AA', 'LightSeaGreen'
    '3C','B3','71', 'MediumSeaGreen'
    '2E','8B','57', 'SeaGreen'
    '8F','BC','8F', 'DarkSeaGreen'
    '22','8B','22', 'ForestGreen'
    '00','64','00', 'DarkGreen'
    '6B','8E','23', 'OliveDrab'
    '80','80','00', 'Olive'
    '55','6B','2F', 'DarkOliveGreen'
    '00','80','80', 'Teal'
    %Blue colors
    '00','00','FF', 'Blue'
    'AD','D8','E6', 'LightBlue'
    'B0','E0','E6', 'PowderBlue'
    'AF','EE','EE', 'PaleTurquoise'
    '40','E0','D0', 'Turquoise'
    '48','D1','CC', 'MediumTurquoise'
    '00','CE','D1', 'DarkTurquoise'
    'E0','FF','FF', 'LightCyan'
    '00','FF','FF', 'Cyan'
    '00','FF','FF', 'Aqua'
    '00','8B','8B', 'DarkCyan'
    '5F','9E','A0', 'CadetBlue'
    'B0','C4','DE', 'LightSteelBlue'
    '46','82','B4', 'SteelBlue'
    '87','CE','FA', 'LightSkyBlue'
    '87','CE','EB', 'SkyBlue'
    '00','BF','FF', 'DeepSkyBlue'
    '1E','90','FF', 'DodgerBlue'
    '64','95','ED', 'CornflowerBlue'
    '41','69','E1', 'RoyalBlue'
    '00','00','CD', 'MediumBlue'
    '00','00','8B', 'DarkBlue'
    '00','00','80', 'Navy'
    '19','19','70', 'MidnightBlue'
    %Purple colors
    '80','00','80', 'Purple'
    'E6','E6','FA', 'Lavender'
    'D8','BF','D8', 'Thistle'
    'DD','A0','DD', 'Plum'
    'EE','82','EE', 'Violet'
    'DA','70','D6', 'Orchid'
    'FF','00','FF', 'Fuchsia'
    'FF','00','FF', 'Magenta'
    'BA','55','D3', 'MediumOrchid'
    '93','70','DB', 'MediumPurple'
    '99','66','CC', 'Amethyst'
    '8A','2B','E2', 'BlueViolet'
    '94','00','D3', 'DarkViolet'
    '99','32','CC', 'DarkOrchid'
    '8B','00','8B', 'DarkMagenta'
    '6A','5A','CD', 'SlateBlue'
    '48','3D','8B', 'DarkSlateBlue'
    '7B','68','EE', 'MediumSlateBlue'
    '4B','00','82', 'Indigo'
    %Gray repeated with spelling grey
    '80','80','80', 'Grey'
    'D3','D3','D3', 'LightGrey'
    'A9','A9','A9', 'DarkGrey'
    '69','69','69', 'DimGrey'
    '77','88','99', 'LightSlateGrey'
    '70','80','90', 'SlateGrey'
    '2F','4F','4F', 'DarkSlateGrey'
    };
  hex = css(:,1:3);
  name = css(:,4);
end
```
