## Supplementary material for "An innovative optical context to make honeybees crash repeatedly": Data set of honeybees flights: script1.m

```
% Programme traitement des données pour ICIV2019
% Retravailler par Julien Serres

clc; clear all; close all
load('trajectoires.mat')

%-------------------------- CONTROLE MIROIR BAS ----------------------------

% Trajectoires en condition contrôle avec les abeilles du groupe demi miroir bas : f8 (g)
figure(1)
subplot(2,1,1);
% Moyennes de chaques trajectoires (utiles pour faire les boîtes à
% moustaches)
z_CC_DMB = [];
z_CC1 = [];
z_CC2 = [];

taille_ligne= 1.5; 


% Tracé des trajectoires :
% plot([CC_B59.x]*128/139,[CC_B59.z]*9/12,'color',[0.2 0.2 0.4])
% z_CC_DMB = [z_CC_DMB mean([CC_B59.z]*9/12)];
% [z_CC_DMB_z1,z_CC_DMB_z2]=Z_CC_DMB2([CC_B59.x]*128/139,[CC_B59.z]*9/12);
% z_CC_DMB_p1 = [z_CC_DMB_p1 mean(z_CC_DMB_z1)];
% z_CC_DMB_p2 = [z_CC_DMB_p2 mean(z_CC_DMB_z2)];
% m=size(CC_B59,2);
% ToF_CC_DMB = [ToF_CC_DMB (1.3*((m)/100))/([CC_B59(1,m).x]*128/139)];
% [ToF_CC(n+1),TbC_MB(n+1)]=Time_of_Flight_DMB([CC_B59.x]*128/139,[CC_B59.z]*9/12,SeuilZ);
% n=n+1;
% hold on
plot([CC_B60.x]*130/max([CC_B60.x]*100),[CC_B60.z],'color',[0.5 0.5 0.5],'LineWidth',taille_ligne)
%plot([CC_B60.x]*130/max([CC_B60.x]*100),[CC_B60.z],'color',[0.2 0.2 0.6],'LineWidth',taille_ligne)
%plot([CC_B60.x]*118/131,[CC_B60.z]*9/13,'color',[0.2 0.2 0.6],'LineWidth',taille_ligne)

CCZ60 = [CC_B60.z];
CCX60 = [CC_B60.x]*130/(max([CC_B60.x]*100));
Index_CC1 = find(CCX60<=0.62);
Index_CC2 = Index_CC1:length(CCX60);

z_CC1 = [z_CC1 mean(CCZ60(Index_CC1))];
z_CC2 = [z_CC2 mean(CCZ60(Index_CC2))];
clear Index_CC1 Index_CC2

hold on
%plot([CC_B61.x]*130/max([CC_B61.x]*100),[CC_B61.z],'color',[0.2 0.2 0.8],'LineWidth',taille_ligne)
plot([CC_B61.x]*130/max([CC_B61.x]*100),[CC_B61.z],'color',[0.5 0.5 0.5],'LineWidth',taille_ligne)
%plot([CC_B61.x]*128/141,[CC_B61.z]*9/13,'color',[0.2 0.2 0.8],'LineWidth',taille_ligne)

CCZ61 = [CC_B61.z];
CCX61 = [CC_B61.x]*130/(max([CC_B61.x]*100));
Index_CC1 = find(CCX61<=0.62);
Index_CC2 = Index_CC1:length(CCX61);

z_CC1 = [z_CC1 mean(CCZ61(Index_CC1))];
z_CC2 = [z_CC2 mean(CCZ61(Index_CC2))];
clear Index_CC1 Index_CC2

hold on
plot([CC_B62.x]*130/max([CC_B62.x]*100),[CC_B62.z],'color',[0.5 0.5 0.5],'LineWidth',taille_ligne)
%plot([CC_B62.x]*130/max([CC_B62.x]*100),[CC_B62.z],'color',[0.2 0.4 0.2],'LineWidth',taille_ligne)
%plot([CC_B62.x]*128/141,[CC_B62.z]*11/14,'color',[0.2 0.4 0.2],'LineWidth',taille_ligne)

CCZ62 = [CC_B62.z];
CCX62 = [CC_B62.x]*130/(max([CC_B62.x]*100));
Index_CC1 = find(CCX62<=0.62);
Index_CC2 = Index_CC1:length(CCX62);

z_CC1 = [z_CC1 mean(CCZ62(Index_CC1))];
z_CC2 = [z_CC2 mean(CCZ62(Index_CC2))];
clear Index_CC1 Index_CC2

hold on
%plot([CC_B63.x]*130/max([CC_B63.x]*100),[CC_B63.z],'color',[0.2 0.4 0.4],'LineWidth',taille_ligne)
plot([CC_B63.x]*130/max([CC_B63.x]*100),[CC_B63.z],'color',[0.5 0.5 0.5],'LineWidth',taille_ligne)
%plot([CC_B63.x]*128/140,[CC_B63.z]*11/14,'color',[0.2 0.4 0.4],'LineWidth',taille_ligne)

CCZ63 = [CC_B63.z];
CCX63 = [CC_B63.x]*130/(max([CC_B63.x]*100));
Index_CC1 = find(CCX63<=0.62);
Index_CC2 = Index_CC1:length(CCX63);

z_CC1 = [z_CC1 mean(CCZ63(Index_CC1))];
z_CC2 = [z_CC2 mean(CCZ63(Index_CC2))];
clear Index_CC1 Index_CC2

hold on
%plot([CC_B64.x]*130/max([CC_B64.x]*100),[CC_B64.z],'color',[0.2 0.4 0.6],'LineWidth',taille_ligne)
plot([CC_B64.x]*130/max([CC_B64.x]*100),[CC_B64.z],'color',[0.5 0.5 0.5],'LineWidth',taille_ligne)
%plot([CC_B64.x]*128/141,[CC_B64.z]*11/14,'color',[0.2 0.4 0.6],'LineWidth',taille_ligne)

CCZ64 = [CC_B64.z];
CCX64 = [CC_B64.x]*130/(max([CC_B64.x]*100));
Index_CC1 = find(CCX64<=0.62);
Index_CC2 = Index_CC1:length(CCX64);

z_CC1 = [z_CC1 mean(CCZ64(Index_CC1))];
z_CC2 = [z_CC2 mean(CCZ64(Index_CC2))];
clear Index_CC1 Index_CC2

hold on
%plot([CC_B65.x]*130/max([CC_B65.x]*100),[CC_B65.z],'color',[0.2 0.4 0.8],'LineWidth',taille_ligne)
plot([CC_B65.x]*130/max([CC_B65.x]*100),[CC_B65.z],'color',[0.5 0.5 0.5],'LineWidth',taille_ligne)
%plot([CC_B65.x]*128/114,[CC_B65.z]*9/13,'color',[0.2 0.4 0.8],'LineWidth',taille_ligne)

CCZ65 = [CC_B65.z];
CCX65 = [CC_B65.x]*130/(max([CC_B65.x]*100));
Index_CC1 = find(CCX65<=0.62);
Index_CC2 = Index_CC1:length(CCX65);

z_CC1 = [z_CC1 mean(CCZ65(Index_CC1))];
z_CC2 = [z_CC2 mean(CCZ65(Index_CC2))];
clear Index_CC1 Index_CC2

hold on
%plot([CC_B66.x]*130/max([CC_B66.x]*100),[CC_B66.z],'color',[0.2 0.6 0.2],'LineWidth',taille_ligne);
plot([CC_B66.x]*130/max([CC_B66.x]*100),[CC_B66.z],'color',[0.5 0.5 0.5],'LineWidth',taille_ligne);
%plot([CC_B66.x]*126/140,[CC_B66.z]*11/14,'color',[0.2 0.6 0.2],'LineWidth',taille_ligne);

CCZ65 = [CC_B65.z];
CCX65 = [CC_B65.x]*130/(max([CC_B65.x]*100));
Index_CC1 = find(CCX65<=0.62);
Index_CC2 = Index_CC1:length(CCX65);

z_CC1 = [z_CC1 mean(CCZ65(Index_CC1))];
z_CC2 = [z_CC2 mean(CCZ65(Index_CC2))];
clear Index_CC1 Index_CC2

hold on
%plot([CC_B67.x]*130/max([CC_B67.x]*100),[CC_B67.z],'color',[0.2 0.6 0.4],'LineWidth',taille_ligne)
plot([CC_B67.x]*130/max([CC_B67.x]*100),[CC_B67.z],'color',[0.5 0.5 0.5],'LineWidth',taille_ligne)
%plot([CC_B67.x]*128/141,[CC_B67.z]*11/14,'color',[0.2 0.6 0.4],'LineWidth',taille_ligne)

CCZ67 = [CC_B67.z];
CCX67 = [CC_B67.x]*130/(max([CC_B67.x]*100));
Index_CC1 = find(CCX67<=0.62);
Index_CC2 = Index_CC1:length(CCX67);

z_CC1 = [z_CC1 mean(CCZ67(Index_CC1))];
z_CC2 = [z_CC2 mean(CCZ67(Index_CC2))];
clear Index_CC1 Index_CC2


hold on
%plot([CC_B68.x]*130/max([CC_B68.x]*100),[CC_B68.z],'color',[0.2 0.6 0.6],'LineWidth',taille_ligne);
plot([CC_B68.x]*130/max([CC_B68.x]*100),[CC_B68.z],'color',[0.5 0.5 0.5],'LineWidth',taille_ligne);
%plot([CC_B68.x]*129/140,[CC_B68.z]*9/11,'color',[0.2 0.6 0.6],'LineWidth',taille_ligne);

CCZ68 = [CC_B68.z];
CCX68 = [CC_B68.x]*130/(max([CC_B68.x]*100));
Index_CC1 = find(CCX68<=0.62);
Index_CC2 = Index_CC1:length(CCX68);

z_CC1 = [z_CC1 mean(CCZ68(Index_CC1))];
z_CC2 = [z_CC2 mean(CCZ68(Index_CC2))];
clear Index_CC1 Index_CC2

hold on
%plot([CC_B69.x]*130/max([CC_B69.x]*100),[CC_B69.z],'color',[0.2 0.6 0.8],'LineWidth',taille_ligne)
plot([CC_B69.x]*130/max([CC_B69.x]*100),[CC_B69.z],'color',[0.5 0.5 0.5],'LineWidth',taille_ligne)
%plot([CC_B69.x]*128/140,[CC_B69.z]*7/8,'color',[0.2 0.6 0.8],'LineWidth',taille_ligne)
CCZ69 = [CC_B69.z];
CCX69 = [CC_B69.x]*130/(max([CC_B69.x]*100));
Index_CC1 = find(CCX69<=0.62);
Index_CC2 = Index_CC1:length(CCX69);

z_CC1 = [z_CC1 mean(CCZ69(Index_CC1))];
z_CC2 = [z_CC2 mean(CCZ69(Index_CC2))];
clear Index_CC1 Index_CC2

% hold on
% plot([CC_B70.x]*128/140,[CC_B70.z]*11/13,'color',[0.2 0.8 0.2])
% z_CC_DMB = [z_CC_DMB mean([CC_B70.z]*11/13)];
% [z_CC_DMB_z1,z_CC_DMB_z2]=Z_CC_DMB_2([CC_B70.x]*128/140,[CC_B70.z]*11/13);
% z_CC_DMB_p1 = [z_CC_DMB_p1 mean(z_CC_DMB_z1)];
% z_CC_DMB_p2 = [z_CC_DMB_p2 mean(z_CC_DMB_z2)];
% m=size(CC_B70,2);
% ToF_CC_DMB = [ToF_CC_DMB (1.3*((m)/100))/([CC_B70(1,m).x]*128/140)];
% [ToF_CC(n+1),TbC_MB(n+1)]=Time_of_Flight_DMB([CC_B70.x]*128/140,[CC_B70.z]*11/13,SeuilZ);
% n=n+1;
hold on
%plot([CC_B72.x]*130/max([CC_B72.x]*100),[CC_B72.z],'color',[0.2 0.8 0.4],'LineWidth',taille_ligne)
plot([CC_B72.x]*130/max([CC_B72.x]*100),[CC_B72.z],'color',[0.5 0.5 0.5],'LineWidth',taille_ligne)
%plot([CC_B72.x]*126/140,[CC_B72.z]*11/13,'color',[0.2 0.8 0.4],'LineWidth',taille_ligne)
CCZ72 = [CC_B72.z];
CCX72 = [CC_B72.x]*130/(max([CC_B72.x]*100));
Index_CC1 = find(CCX72<=0.62);
Index_CC2 = Index_CC1:length(CCX72);

z_CC1 = [z_CC1 mean(CCZ72(Index_CC1))];
z_CC2 = [z_CC2 mean(CCZ72(Index_CC2))];
clear Index_CC1 Index_CC2

hold on
%plot([CC_B73.x]*126/141,[CC_B73.z]*12/15,'color',[0.2 0.8 0.6],'LineWidth',taille_ligne)
plot([CC_B73.x]*130/max([CC_B73.x]),[CC_B73.z],'color',[0.5 0.5 0.5],'LineWidth',taille_ligne)

CCZ73 = [CC_B73.z];
CCX73 = [CC_B73.x]*130/(max([CC_B73.x]*100));
Index_CC1 = find(CCX73<=0.62);
Index_CC2 = Index_CC1:length(CCX73);

z_CC1 = [z_CC1 mean(CCZ73(Index_CC1))];
z_CC2 = [z_CC2 mean(CCZ73(Index_CC2))];
clear Index_CC1 Index_CC2

hold on
%plot([CC_B74.x]*130/max([CC_B74.x]*100),[CC_B74.z],'color',[0.2 0.8 0.8],'LineWidth',taille_ligne)
plot([CC_B74.x]*130/max([CC_B74.x]*100),[CC_B74.z],'color',[0.5 0.5 0.5],'LineWidth',taille_ligne)
%plot([CC_B74.x]*126/93,[CC_B74.z]*9/12,'color',[0.2 0.8 0.8],'LineWidth',taille_ligne)

CCZ74 = [CC_B74.z];
CCX74 = [CC_B74.x]*130/(max([CC_B74.x]*100));
Index_CC1 = find(CCX74<=0.62);
Index_CC2 = Index_CC1:length(CCX74);

z_CC1 = [z_CC1 mean(CCZ74(Index_CC1))];
z_CC2 = [z_CC2 mean(CCZ74(Index_CC2))];
clear Index_CC1 Index_CC2

hold on
%plot([CC_B75.x]*130/max([CC_B75.x]*100),[CC_B75.z],'color',[0.4 0.2 0.4],'LineWidth',taille_ligne)
plot([CC_B75.x]*130/max([CC_B75.x]*100),[CC_B75.z],'color',[0.5 0.5 0.5],'LineWidth',taille_ligne)
%plot([CC_B75.x]*126/140,[CC_B75.z]*11/14,'color',[0.4 0.2 0.4],'LineWidth',taille_ligne)

CCZ75 = [CC_B75.z];
CCX75 = [CC_B75.x]*130/(max([CC_B75.x]*100));
Index_CC1 = find(CCX75<=0.62);
Index_CC2 = Index_CC1:length(CCX75);

z_CC1 = [z_CC1 mean(CCZ75(Index_CC1))];
z_CC2 = [z_CC2 mean(CCZ75(Index_CC2))];
clear Index_CC1 Index_CC2

hold on
%plot([CC_B78.x]*130/max([CC_B78.x]*100),[CC_B78.z],'color',[0.4 0.2 0.8],'LineWidth',taille_ligne)
plot([CC_B78.x]*130/max([CC_B78.x]*100),[CC_B78.z],'color',[0.5 0.5 0.5],'LineWidth',taille_ligne)
%plot([CC_B78.x]*124/155,[CC_B78.z]*8/22,'color',[0.4 0.2 0.8],'LineWidth',taille_ligne)

CCZ78 = [CC_B78.z];
CCX78 = [CC_B78.x]*130/(max([CC_B78.x]*100));
Index_CC1 = find(CCX78<=0.62);
Index_CC2 = Index_CC1:length(CCX78);

z_CC1 = [z_CC1 mean(CCZ78(Index_CC1))];
z_CC2 = [z_CC2 mean(CCZ78(Index_CC2))];
clear Index_CC1 Index_CC2

hold on
plot([0.62 0.62],[0 0.71],'color',[0 0 0],'LineStyle','--')
% Mise en forme du graphique :
% xlabel('Distance travelled (m)')
% ylabel('Altitude (m)')
xticks([0 0.62 1.3]) % On programme l'échelle à afficher en x
yticks([-0.02 0 0.09 0.14 0.71]) % On programme l'échelle à afficher en y
xticklabels({'0','0.62','1.3'}) % On programme le texte à écrire sous l'échelle en x
yticklabels({'','0','0.09','0.14','0.71'}) % On programme le texte à écrire sous l'échelle en y
axis([0 1.3 -0.02 0.73]) % On donne la limite des axes
text(0.1,0.6,'(a) Control condition') % On demande à écrire "(g)" en haut à gauche dans le graphique
n = 15;
str={n};
text(0.7,0.6,'n = ');
text(0.77,0.6,str);
ylabel('Altitude (m)')
% t=0.01*round(median(ToF_CC_DMB)*100);
% text(1,0.55,strcat('ToF =',num2str(t),'  s'));
%xlabel('Distance travelled (m)')

% Dessin des motifs et miroirs dans le graphique
offsety = 7.1; % 100 mm

% 2 épaisseur

for p = 1 : 22
        m = (p-0.01)*0.1-0.1 ;
        rectangle('Position',[m,offsety/10,3,0.2],'FaceColor','w','EdgeColor',[1 0 0]);   %
        rectangle('Position',[0.03+m,offsety/10,1,0.2],'FaceColor','r','EdgeColor',[1 0 0]); %
        rectangle('Position',[0.04+m,offsety/10,1,0.2],'FaceColor','w','EdgeColor',[1 0 0]); % 11 periode plafond
        rectangle('Position',[0.05+m,offsety/10,1,0.2],'FaceColor','r','EdgeColor',[1 0 0]); %
        rectangle('Position',[0.06+m,offsety/10,1,0.2],'FaceColor','w','EdgeColor',[1 0 0]); %
        rectangle('Position',[0.07+m,offsety/10,3,0.2],'FaceColor','r','EdgeColor',[1 0 0]); %
end

for p = 1 : 22
        m = (p-0.01)*0.1-0.1 ;
        rectangle('Position',[m,-0.2,3,0.2],'FaceColor','w','EdgeColor',[1 0 0]);   %
        rectangle('Position',[0.03+m,-0.2,1,0.2],'FaceColor','r','EdgeColor',[1 0 0]); %
        rectangle('Position',[0.04+m,-0.2,1,0.2],'FaceColor','w','EdgeColor',[1 0 0]); % 11 periode plafond
        rectangle('Position',[0.05+m,-0.2,1,0.2],'FaceColor','r','EdgeColor',[1 0 0]); %
        rectangle('Position',[0.06+m,-0.2,1,0.2],'FaceColor','w','EdgeColor',[1 0 0]); %
        rectangle('Position',[0.07+m,-0.2,3,0.2],'FaceColor','r','EdgeColor',[1 0 0]); %
end

title('16th flight recorded in control condition after training','FontWeight','normal')
text(0.2,0.5,'z = 9.4 \pm 2.9 cm')
text(0.7,0.5,'z = 11.6 \pm 2.7 cm')

'z_CC1'
mean(z_CC1)
std(z_CC1)

'z_CC2'
mean(z_CC2)
std(z_CC2)

%--------------------------- DEMI MIROIR BAS --------------------------------------
% Trajectoires en condition Demi Miroir Bas (DMB) : f9 (h)

subplot(2,1,2);
% Moyennes de chaques trajectoires (utiles pour faire les boîtes à moustaches)
z_DMB = [];

z_DMB_CC = [];
z_DMB_MB = [];

n=0;
SeuilZ = 30; % seuil de vol à 30 mm

%Tracé des trajectoires :
hold on
Z60 = [DMB_B60.z];
%Z60 = ([DMB_B60.z]-Z60(181))/(0.71-2*Z60(181));
X60 = [DMB_B60.x]*130/(max([DMB_B66.x]*100));
%plot(X60(1:182),[Z60(1:181) 0],'color',[0.2 0.2 0.6],'LineWidth',taille_ligne)
plot(X60(1:182),[Z60(1:181) 0],'color',[0.5 0.5 0.5],'LineWidth',taille_ligne)
%plot([DMB_B60.x]*97/116,[DMB_B60.z]*9/11,'color',[0.2 0.2 0.6])
Index_DMB_CC = find(X60<=0.62);
Index_DMB_MB = Index_DMB_CC:181;

z_DMB_CC = [z_DMB_CC mean(Z60(Index_DMB_CC))];
z_DMB_MB = [z_DMB_MB mean(Z60(Index_DMB_MB))];
clear Index_DMB_CC Index_DMB_MB

hold on
Z61 = [DMB_B61.z];
%Z61 = ([DMB_B61.z]-Z61(162))/(0.71-2*Z61(162));
X61 = [DMB_B61.x]*130/(max([DMB_B61.x]*100));
%plot(X61(1:163),[Z61(1:162) 0],'color',[0.2 0.2 0.8],'LineWidth',taille_ligne)
plot(X61(1:163),[Z61(1:162) 0],'color',[0.5 0.5 0.5],'LineWidth',taille_ligne)

%plot([DMB_B61.x]*125/138,[DMB_B61.z]*8/12,'color',[0.2 0.2 0.8])
Index_DMB_CC = find(X61<=0.62);
Index_DMB_MB = Index_DMB_CC:162;

z_DMB_CC = [z_DMB_CC mean(Z61(Index_DMB_CC))];
z_DMB_MB = [z_DMB_MB mean(Z61(Index_DMB_MB))];
clear Index_DMB_CC Index_DMB_MB

hold on
Z62 = [DMB_B62.z];
%Z62 = ([DMB_B62.z]-Z62(244))/(0.71-2*Z62(244));
X62 = [DMB_B62.x];
%plot(X62(1:245),[Z62(1:244) 0],'color',[0.2 0.4 0.2],'LineWidth',taille_ligne)
% Celle-ci ne vole pas au-dessus de la bâche
plot(X62(1:245),[Z62(1:244) 0],'color',[0.5 0.5 0.5],'LineWidth',taille_ligne)
%plot([DMB_B62.x]*77/93,[DMB_B62.z]*9/11,'color',[0.2 0.4 0.2])
Index_DMB_CC = find(X62<=0.62);
Index_DMB_MB = 0; % le z au delà est zéro

z_DMB_CC = [z_DMB_CC mean(Z62(Index_DMB_CC))];
z_DMB_MB = [z_DMB_MB 0];
clear Index_DMB_CC Index_DMB_MB


hold on
Z63 = [DMB_B63.z];
%Z63 = ([DMB_B63.z]-Z63(185))/(0.71-2*Z63(185));
X63 = [DMB_B63.x];
%plot(X63(1:186),[Z63(1:185) 0],'color',[0.2 0.4 0.4],'LineWidth',taille_ligne)
plot(X63(1:186),[Z63(1:185) 0],'color',[0.5 0.5 0.5],'LineWidth',taille_ligne)
%plot([DMB_B63.x]*106/122,[DMB_B63.z]*13/9,'color',[0.2 0.4 0.4])
Index_DMB_CC = find(X63<=0.62);
Index_DMB_MB = Index_DMB_CC:185;

z_DMB_CC = [z_DMB_CC mean(Z63(Index_DMB_CC))];
z_DMB_MB = [z_DMB_MB mean(Z63(Index_DMB_MB))];
clear Index_DMB_CC Index_DMB_MB

hold on
Z65 = [DMB_B65.z];
%Z65 = ([DMB_B65.z]-Z65(109))/(0.71-2*Z65(109));
X65 = [DMB_B65.x];
%plot(X65(1:110),[Z65(1:109) 0],'color',[0.2 0.4 0.8],'LineWidth',taille_ligne)
plot(X65(1:110),[Z65(1:109) 0],'color',[0.5 0.5 0.5],'LineWidth',taille_ligne)
%plot([DMB_B65.x]*128/96,[DMB_B65.z]*9/12,'color',[0.2 0.4 0.8])
Index_DMB_CC = find(X65<=0.62);
Index_DMB_MB = Index_DMB_CC:109;

z_DMB_CC = [z_DMB_CC mean(Z65(Index_DMB_CC))];
z_DMB_MB = [z_DMB_MB mean(Z65(Index_DMB_MB))];
clear Index_DMB_CC Index_DMB_MB

hold on
Z66 = [DMB_B66.z];
%Z66 = ([DMB_B66.z]-Z66(161))/(0.71-2*Z66(161));
X66 = [DMB_B66.x]*130/(max([DMB_B66.x]*100));
%plot(X66(1:162),[Z66(1:161) 0],'color',[0.2 0.6 0.2],'LineWidth',taille_ligne)
plot(X66(1:162),[Z66(1:161) 0],'color',[0.5 0.5 0.5],'LineWidth',taille_ligne)
%plot([DMB_B66.x]*130/140,[DMB_B66.z]*10/14,'color',[0.2 0.6 0.2])
Index_DMB_CC = find(X66<=0.62);
Index_DMB_MB = Index_DMB_CC:161;

z_DMB_CC = [z_DMB_CC mean(Z66(Index_DMB_CC))];
z_DMB_MB = [z_DMB_MB mean(Z66(Index_DMB_MB))];
%clear Index_DMB_CC Index_DMB_MB

hold on
Z67 = [DMB_B67.z];
%Z67 = ([DMB_B67.z]-Z67(96))/(0.71-2*Z67(96));
X67 = [DMB_B67.x]*130/(max([DMB_B67.x]*100));
%plot(X67(1:97),[Z67(1:96) 0],'color',[0.2 0.6 0.4],'LineWidth',taille_ligne)
plot(X67(1:97),[Z67(1:96) 0],'color',[0.5 0.5 0.5],'LineWidth',taille_ligne)
%plot([DMB_B67.x]*129/140,[DMB_B67.z]*9/12,'color',[0.2 0.6 0.4])
Index_DMB_CC = find(X67<=0.62);
Index_DMB_MB = Index_DMB_CC:96;

z_DMB_CC = [z_DMB_CC mean(Z67(Index_DMB_CC))];
z_DMB_MB = [z_DMB_MB mean(Z67(Index_DMB_MB))];
clear Index_DMB_CC Index_DMB_MB

hold on
Z68 = [DMB_B68.z];
%Z68 = ([DMB_B68.z]-Z68(309))/(0.71-2*Z68(309));
X68 = [DMB_B68.x];
%plot(X68(1:310),[Z68(1:309) 0],'color',[0.2 0.6 0.6],'LineWidth',taille_ligne)
plot(X68(1:310),[Z68(1:309) 0],'color',[0.5 0.5 0.5],'LineWidth',taille_ligne)
%plot([DMB_B68.x]*66/84,[DMB_B68.z]*9/12,'color',[0.2 0.6 0.6])
Index_DMB_CC = find(X68<=0.62);
Index_DMB_MB = Index_DMB_CC:309;

z_DMB_CC = [z_DMB_CC mean(Z68(Index_DMB_CC))];
z_DMB_MB = [z_DMB_MB mean(Z68(Index_DMB_MB))];
clear Index_DMB_CC Index_DMB_MB
%return

hold on
Z69 = [DMB_B69.z];
%Z69 = ([DMB_B69.z]-Z69(204))/(0.71-2*Z69(204));
X69 = [DMB_B69.x]*130/(max([DMB_B69.x]*100));
%plot(X69(1:205),[Z69(1:204) 0],'color',[0.2 0.6 0.8],'LineWidth',taille_ligne)
plot(X69(1:205),[Z69(1:204) 0],'color',[0.5 0.5 0.5],'LineWidth',taille_ligne)
%plot([DMB_B69.x]*130/141,[DMB_B69.z]*9/13,'color',[0.2 0.6 0.8])
Index_DMB_CC = find(X69<=0.62);
Index_DMB_MB = Index_DMB_CC:204;

z_DMB_CC = [z_DMB_CC mean(Z69(Index_DMB_CC))];
z_DMB_MB = [z_DMB_MB mean(Z69(Index_DMB_MB))];
clear Index_DMB_CC Index_DMB_MB


hold on
Z70 = [DMB_B70.z];
X70 = [DMB_B70.x];
%plot(X70(1:end),Z70(1:end),'color',[0.2 0.8 0.2],'LineWidth',taille_ligne)
plot(X70(1:end),Z70(1:end),'color',[0.5 0.5 0.5],'LineWidth',taille_ligne)
% ne finie pas au sol
%plot([DMB_B70.x]*124/97,[DMB_B70.z]*9/12,'color',[0.2 0.8 0.2])
Index_DMB_CC = find(X70<=0.62);
Index_DMB_MB = Index_DMB_CC:length(X70);

z_DMB_CC = [z_DMB_CC mean(Z70(Index_DMB_CC))];
z_DMB_MB = [z_DMB_MB mean(Z70(Index_DMB_MB))];
%clear Index_DMB_CC Index_DMB_MB

hold on
Z72 = [DMB_B72.z];
%Z72 = ([DMB_B72.z]-Z72(175))/(0.71-2*Z72(175));
X72 = [DMB_B72.x]*130/(max([DMB_B72.x]*100));
%plot(X72(1:176),[Z72(1:175) 0],'color',[0.2 0.8 0.4],'LineWidth',taille_ligne)
plot(X72(1:176),[Z72(1:175) 0],'color',[0.5 0.5 0.5],'LineWidth',taille_ligne)
%plot([DMB_B72.x]*128/140,[DMB_B72.z]*9/12,'color',[0.2 0.8 0.4])
Index_DMB_CC = find(X72<=0.62);
Index_DMB_MB = Index_DMB_CC:175;

z_DMB_CC = [z_DMB_CC mean(Z72(Index_DMB_CC))];
z_DMB_MB = [z_DMB_MB mean(Z72(Index_DMB_MB))];
clear Index_DMB_CC Index_DMB_MB

hold on
Z73 = [DMB_B73.z];
%Z73 = ([DMB_B73.z]-Z73(131))/(0.71-2*Z73(131));
X73 = [DMB_B73.x]*130/(max([DMB_B72.x]*100));
%plot(X73(1:132),[Z73(1:131) 0],'color',[0.2 0.8 0.6],'LineWidth',taille_ligne)
plot(X73(1:132),[Z73(1:131) 0],'color',[0.5 0.5 0.5],'LineWidth',taille_ligne)
%plot([DMB_B73.x]*123/124,[DMB_B73.z]*9/12,'color',[0.2 0.8 0.6])
Index_DMB_CC = find(X73<=0.62);
Index_DMB_MB = Index_DMB_CC:131;

z_DMB_CC = [z_DMB_CC mean(Z73(Index_DMB_CC))];
z_DMB_MB = [z_DMB_MB mean(Z73(Index_DMB_MB))];
clear Index_DMB_CC Index_DMB_MB

hold on
Z74 = [DMB_B74.z];
%Z74 = ([DMB_B74.z]-Z74(152))/(0.71-2*Z74(152));
X74 = [DMB_B74.x]*130/(max([DMB_B74.x]*100));
%plot(X74(1:153),[Z74(1:152) 0],'color',[0.2 0.8 0.8],'LineWidth',taille_ligne)
plot(X74(1:153),[Z74(1:152) 0],'color',[0.5 0.5 0.5],'LineWidth',taille_ligne)
%plot([DMB_B74.x]*130/140,[DMB_B74.z]*9/13,'color',[0.2 0.8 0.8])
Index_DMB_CC = find(X74<=0.62);
Index_DMB_MB = Index_DMB_CC:133;

z_DMB_CC = [z_DMB_CC mean(Z74(Index_DMB_CC))];
z_DMB_MB = [z_DMB_MB mean(Z74(Index_DMB_MB))];
clear Index_DMB_CC Index_DMB_MB

hold on
Z75 = [DMB_B75.z];
%Z75 = ([DMB_B75.z]-Z75(154))/(0.71-2*Z75(154));
X75 = [DMB_B75.x];
%plot(X75(1:155),[Z75(1:154) 0],'color',[0.4 0.2 0.4],'LineWidth',taille_ligne)
plot(X75(1:155),[Z75(1:154) 0],'color',[0.5 0.5 0.5],'LineWidth',taille_ligne)
%plot([DMB_B75.x]*96/109,[DMB_B75.z]*9/12,'color',[0.4 0.2 0.4])
Index_DMB_CC = find(X75<=0.62);
Index_DMB_MB = Index_DMB_CC:154;

z_DMB_CC = [z_DMB_CC mean(Z75(Index_DMB_CC))];
z_DMB_MB = [z_DMB_MB mean(Z75(Index_DMB_MB))];
clear Index_DMB_CC Index_DMB_MB

hold on
Z78 = [DMB_B78.z];
%Z78 = ([DMB_B78.z]-Z78(108))/(0.71-2*Z78(108));
%X78 = [DMB_B78.x]*130/(max([DMB_B78.x]*100));
X78 = [DMB_B78.x];
%plot(X78(1:109),[Z78(1:108) 0],'color',[0.4 0.2 0.8],'LineWidth',taille_ligne)
plot(X78(1:109),[Z78(1:108) 0],'color',[0.5 0.5 0.5],'LineWidth',taille_ligne)
%plot([DMB_B78.x]*130/155,[DMB_B78.z]*9/19,'color',[0.4 0.2 0.8])
Index_DMB_CC = find(X78<=0.62);
Index_DMB_MB = Index_DMB_CC:138;

z_DMB_CC = [z_DMB_CC mean(Z78(Index_DMB_CC))];
z_DMB_MB = [z_DMB_MB mean(Z78(Index_DMB_MB))];
clear Index_DMB_CC Index_DMB_MB

hold on
plot([0.62 0.62],[0 0.71],'color',[0 0 0],'LineStyle','--')

% Mise en forme du graphique :
% xlabel('Distance travelled (m)')
% ylabel('Altitude (m)')
xticks([0 0.62 1.3]) % On programme l'échelle à afficher en x
yticks([-0.02 0 0.09 0.14 0.71]) % On programme l'échelle à afficher en y
xticklabels({'0','0.62','1.3'}) % On programme le texte à écrire sous l'échelle en x
yticklabels({'','0','0.09','0.14','0.71'}) % On programme le texte à écrire sous l'échelle en y
axis([0 1.3 -0.02 0.73]) % On donne la limite des axes
text(0.1,0.6,'(b) Half mirror on the floor') % On demande à écrire "(h)" en haut à gauche dans le graphique
text(0.7,0.6,'n = 15')
% TbC_DMB=TbC;
% ToF=0.01*round(median(ToF)*100);
% text(0.1,0.50,strcat('ToF = ',num2str(ToF),'  s'))
% TbC=0.01*round(median(TbC)*100);
% text(1,0.50,strcat('TbC = ',num2str(TbC),'  s'))

% Dessin des motifs et miroirs dans le graphique
offsety = 7.1; % 100 mm

% 2 épaisseur

for p = 1 : 22
        m = (p-0.01)*0.1-0.1 ;
        rectangle('Position',[m,offsety/10,3,0.2],'FaceColor','w','EdgeColor',[1 0 0]);   %
        rectangle('Position',[0.03+m,offsety/10,1,0.2],'FaceColor','r','EdgeColor',[1 0 0]); %
        rectangle('Position',[0.04+m,offsety/10,1,0.2],'FaceColor','w','EdgeColor',[1 0 0]); % 11 periode plafond
        rectangle('Position',[0.05+m,offsety/10,1,0.2],'FaceColor','r','EdgeColor',[1 0 0]); %
        rectangle('Position',[0.06+m,offsety/10,1,0.2],'FaceColor','w','EdgeColor',[1 0 0]); %
        rectangle('Position',[0.07+m,offsety/10,3,0.2],'FaceColor','r','EdgeColor',[1 0 0]); %
end
    
% 110 x 2 = 220 distance max

for p = 1:6.9
        m = (p-0.01)*0.1-0.1;
        rectangle('Position',[m,-0.2,3,0.2],'FaceColor','w','EdgeColor',[1 0 0]);   %
        rectangle('Position',[0.03+m,-0.2,1,0.2],'FaceColor','r','EdgeColor',[1 0 0]); %
        rectangle('Position',[0.04+m,-0.2,1,0.2],'FaceColor','w','EdgeColor',[1 0 0]); % 11 periode plafond
        rectangle('Position',[0.05+m,-0.2,1,0.2],'FaceColor','r','EdgeColor',[1 0 0]); %
        rectangle('Position',[0.06+m,-0.2,1,0.2],'FaceColor','w','EdgeColor',[1 0 0]); %
        rectangle('Position',[0.07+m,-0.2,3,0.2],'FaceColor','r','EdgeColor',[1 0 0]); %
        rectangle('Position',[0.10+m,-0.2,3,0.2],'FaceColor','w','EdgeColor',[1 0 0]);
end

for p = 1 : 55
        m = (p-0.03)*0.04+0.01 + 0.78;
        hold on
       plot([m-0.2 m+0.1 ],[0 -0.2],'-b','LineWidth',1.3)
end
hold on 
plot([0.62 1.3], [0 0],'-b','LineWidth',1.3)
title('1st flight recorded in half mirror on the floor condition','FontWeight','normal')
xlabel('Distance travelled (m)')
ylabel('Altitude (m)')
%[ax1,h1]=suplabel('Distance travelled (m)','x'); 
%[ax2,h2]=suplabel('Altitude (m)','y');
box on


% 1er vol en demi miroir bas DMB
text(0.2,0.5,'z = 10.4 \pm 3.6 cm')

text(0.7,0.5,'z = 10.4 \pm 3.9 cm')
text(0.7,0.4,'x = 29.5 \pm 14.9 cm')

'z_DMB_CC'
mean(z_DMB_CC)
std(z_DMB_CC)

'z_DMB_MB'
mean(z_DMB_MB)
std(z_DMB_MB)

x_DMB = [X60(181) X61(162) X62(244) X63(185) X65(109) X66(161) X67(96) X68(309) X69(204) max(X70) X72(175) X73(131) X74(152) X75(154) X78(108)];
mean(x_DMB)
std(x_DMB)

print -f1 -dpng -r300 DMB

% figure(2)
% boxplot([z_DMB_CC',z_DMB_MB'],'Notch','on','Labels',{'Control','HLM'})
% ylabel('Altitude (m)')
% title('Half Low Mirror (HLM) condition')
% 
% [h,p] = ttest(z_DMB_CC',z_DMB_MB','Alpha',0.05)
% 
% distance_travelled = [X60(181) X61(162) X62(244) X63(185) X65(109) X66(161) X67(96) X68(309) X69(204) X70(end) X72(175) X73(131) X74(152) X75(154) X78(108)];
% [h,p] = lillietest(distance_travelled);
% [h,p]  = ttest(distance_travelled,1.3,'Tail','right')
% 
% [h,p]  = ttest(z_CC1,z_DMB_CC)
% % p = 0.18
% [h,p]  = ttest(z_CC2,z_DMB_MB)
% p = 0.15


%------------------------- CONTROLE DOUBLE MIROIR -----------------------------------------
% Trajectoires en condition contrôle avec les abeilles du groupe double miroir : f4 (e)

figure(2)
subplot(2,1,1);
% Moyennes de chaques trajectoires (utiles pour faire les boîtes à
% moustaches)
z_CC_DM = [];
n = 0;

%Tracé des trajectoires :
plot([CC_B23.x]*130/max([CC_B23.x]*100),[CC_B23.z],'color',[0.5 0.5 0.5],'LineWidth',taille_ligne)
z_CC_DM = [z_CC_DM mean([CC_B23.z])];
n = n+1;
hold on

plot([CC_B24.x]*130/max([CC_B24.x]*100),[CC_B24.z],'color',[0.5 0.5 0.5],'LineWidth',taille_ligne)
z_CC_DM = [z_CC_DM mean([CC_B24.z])];
n = n+1;
hold on

plot([CC_B25.x]*130/max([CC_B25.x]*100),[CC_B25.z],'color',[0.5 0.5 0.5],'LineWidth',taille_ligne)
z_CC_DM = [z_CC_DM mean([CC_B25.z])];
n = n+1;
hold on

plot([CC_B26.x]*130/max([CC_B26.x]*100),[CC_B26.z],'color',[0.5 0.5 0.5],'LineWidth',taille_ligne)
z_CC_DM = [z_CC_DM mean([CC_B26.z])];
n = n+1;
hold on

plot([CC_B27.x]*130/max([CC_B27.x]*100),[CC_B27.z],'color',[0.5 0.5 0.5],'LineWidth',taille_ligne)
z_CC_DM = [z_CC_DM mean([CC_B27.z])];
n = n+1;
hold on

plot([CC_B28.x]*130/max([CC_B28.x]*100),[CC_B28.z],'color',[0.5 0.5 0.5],'LineWidth',taille_ligne)
z_CC_DM = [z_CC_DM mean([CC_B28.z])];
n = n+1;
hold on

plot([CC_B29.x]*130/max([CC_B29.x]*100),[CC_B29.z],'color',[0.5 0.5 0.5],'LineWidth',taille_ligne)
z_CC_DM = [z_CC_DM mean([CC_B29.z])];
n = n+1;
hold on

plot([CC_B30.x]*130/max([CC_B30.x]*100),[CC_B30.z],'color',[0.5 0.5 0.5],'LineWidth',taille_ligne)
z_CC_DM = [z_CC_DM mean([CC_B30.z])];
n = n+1;
hold on

plot([CC_B31.x]*130/max([CC_B31.x]*100),[CC_B31.z],'color',[0.5 0.5 0.5],'LineWidth',taille_ligne)
z_CC_DM = [z_CC_DM mean([CC_B31.z])];
n = n+1;
hold on

plot([CC_B32.x]*130/max([CC_B32.x]*100),[CC_B32.z],'color',[0.5 0.5 0.5],'LineWidth',taille_ligne)
z_CC_DM = [z_CC_DM mean([CC_B32.z])];
n = n+1;
hold on

plot([CC_B33.x]*130/max([CC_B33.x]*100),[CC_B33.z],'color',[0.5 0.5 0.5],'LineWidth',taille_ligne)
z_CC_DM = [z_CC_DM mean([CC_B33.z])];
n = n+1;
hold on

plot([CC_B34.x]*130/max([CC_B34.x]*100),[CC_B34.z],'color',[0.5 0.5 0.5],'LineWidth',taille_ligne)
z_CC_DM = [z_CC_DM mean([CC_B34.z])];
n = n+1;
hold on

plot([CC_B35.x]*130/max([CC_B35.x]*100),[CC_B35.z],'color',[0.5 0.5 0.5],'LineWidth',taille_ligne)
z_CC_DM = [z_CC_DM mean([CC_B35.z])];
n = n+1;
hold on

plot([CC_B37.x]*130/max([CC_B37.x]*100),[CC_B37.z],'color',[0.5 0.5 0.5],'LineWidth',taille_ligne)
z_CC_DM = [z_CC_DM mean([CC_B37.z])];
n = n+1;
hold on

plot([CC_B54.x]*130/max([CC_B54.x]*100),[CC_B54.z],'color',[0.5 0.5 0.5],'LineWidth',taille_ligne)
z_CC_DM = [z_CC_DM mean([CC_B54.z])];
n = n+1;
hold on


xticks([0 1.3]) % On programme l'échelle à afficher en x
yticks([-0.02 0 0.09 0.14 0.71]) % On programme l'échelle à afficher en y
xticklabels({'0','1.3'}) % On programme le texte à écrire sous l'échelle en x
yticklabels({'','0','0.09','0.14','0.71'}) % On programme le texte à écrire sous l'échelle en y
axis([0 1.3 -0.02 0.73]) % On donne la limite des axes
text(0.1,0.6,'(a) Control condition') % On demande à écrire "(e)" en haut à gauche dans le graphique
str={n};
text(0.7,0.6,'n = ')
text(0.77,0.6,str)
%t=0.01*round(median(ToF_CC_DM)*100);
%text(1,0.55,strcat('ToF =',num2str(t),'  s'))

% Dessin des motifs et miroirs dans le graphique
offsety = 7.1; % 100 mm

% 2 épaisseur

for p = 1 : 22
        m = (p-0.01)*0.1-0.1 ;
        rectangle('Position',[m,offsety/10,3,0.2],'FaceColor','w','EdgeColor',[1 0 0]);   %
        rectangle('Position',[0.03+m,offsety/10,1,0.2],'FaceColor','r','EdgeColor',[1 0 0]); %
        rectangle('Position',[0.04+m,offsety/10,1,0.2],'FaceColor','w','EdgeColor',[1 0 0]); % 11 periode plafond
        rectangle('Position',[0.05+m,offsety/10,1,0.2],'FaceColor','r','EdgeColor',[1 0 0]); %
        rectangle('Position',[0.06+m,offsety/10,1,0.2],'FaceColor','w','EdgeColor',[1 0 0]); %
        rectangle('Position',[0.07+m,offsety/10,3,0.2],'FaceColor','r','EdgeColor',[1 0 0]); %
end

for p = 1 : 22
        m = (p-0.01)*0.1-0.1 ;
        rectangle('Position',[m,-0.2,3,0.2],'FaceColor','w','EdgeColor',[1 0 0]);   %
        rectangle('Position',[0.03+m,-0.2,1,0.2],'FaceColor','r','EdgeColor',[1 0 0]); %
        rectangle('Position',[0.04+m,-0.2,1,0.2],'FaceColor','w','EdgeColor',[1 0 0]); % 11 periode plafond
        rectangle('Position',[0.05+m,-0.2,1,0.2],'FaceColor','r','EdgeColor',[1 0 0]); %
        rectangle('Position',[0.06+m,-0.2,1,0.2],'FaceColor','w','EdgeColor',[1 0 0]); %
        rectangle('Position',[0.07+m,-0.2,3,0.2],'FaceColor','r','EdgeColor',[1 0 0]); %
end

ylabel('Altitude (m)')
title('16th flight recorded in control condition after training','FontWeight','normal')
box on
text(0.7,0.5,'z = 9.7 \pm 3.2 cm')
%--------------------------- DOUBLE MIROIR --------------------------------------------
% Trajectoires en condition double miroir : f2

subplot(2,1,2);

% Moyennes de chaques trajectoires (utiles pour faire les boîtes à moustaches)
z_DM = [];
x_DM = [];
n=0;

%Tracé des trajectoires :
DM_B23X = [DM_B23.x];
DM_B23Z = [DM_B23.z];
Xmax = 125; % valeur de X avant premier crash
plot([DM_B23X(1:Xmax) DM_B23X(Xmax)],[DM_B23Z(1:Xmax) 0],'color',[0.5 0.5 0.5],'LineWidth',taille_ligne)
z_DM = [z_DM mean(DM_B23Z(1:Xmax))];
x_DM = [x_DM DM_B23X(Xmax)];
n=n+1;
hold on


DM_B24X = [DM_B24.x];
DM_B24Z = [DM_B24.z];
Xmax = 90; % valeur de X avant premier crash (23)
plot([DM_B24X(1:Xmax) DM_B24X(Xmax)],[DM_B24Z(1:Xmax) 0],'color',[0.5 0.5 0.5],'LineWidth',taille_ligne)
z_DM = [z_DM mean(DM_B24Z(1:Xmax))];
x_DM = [x_DM DM_B24X(Xmax)];
n=n+1;
hold on

DM_B25X = [DM_B25.x];
DM_B25Z = [DM_B25.z];
Xmax = 80; % valeur de X avant premier crash (20)
plot([DM_B25X(1:Xmax) DM_B25X(Xmax)],[DM_B25Z(1:Xmax) 0],'color',[0.5 0.5 0.5],'LineWidth',taille_ligne)
z_DM = [z_DM mean(DM_B25Z(1:Xmax))];
x_DM = [x_DM DM_B25X(Xmax)];
n=n+1;
hold on

DM_B26X = [DM_B26.x];
DM_B26Z = [DM_B26.z];
Xmax = 28; % valeur de X avant premier crash
plot([DM_B26X(1:Xmax) DM_B26X(Xmax)],[DM_B26Z(1:Xmax) 0],'color',[0.5 0.5 0.5],'LineWidth',taille_ligne)
z_DM = [z_DM mean(DM_B26Z(1:Xmax))];
x_DM = [x_DM DM_B26X(Xmax)];
n=n+1;
hold on

DM_B27X = [DM_B27.x];
DM_B27Z = [DM_B27.z];
Xmax = find(DM_B27Z==min(DM_B27Z)); % valeur de X avant premier crash
plot([DM_B27X(1:Xmax) DM_B27X(Xmax)],[DM_B27Z(1:Xmax) 0],'color',[0.5 0.5 0.5],'LineWidth',taille_ligne)
z_DM = [z_DM mean(DM_B27Z(1:Xmax))];
x_DM = [x_DM DM_B27X(Xmax)];
n=n+1;
hold on

DM_B28X = [DM_B28.x];
DM_B28Z = [DM_B28.z];
Xmax = find(DM_B28Z==min(DM_B28Z)); % valeur de X avant premier crash
plot([DM_B28X(1:Xmax) DM_B28X(Xmax)],[DM_B28Z(1:Xmax) 0],'color',[0.5 0.5 0.5],'LineWidth',taille_ligne)
z_DM = [z_DM mean(DM_B28Z(1:Xmax))];
x_DM = [x_DM DM_B28X(Xmax)];
n=n+1;
hold on

DM_B29X = [DM_B29.x];
DM_B29Z = [DM_B29.z];
Xmax = find(DM_B29Z==min(DM_B29Z)); % valeur de X avant premier crash
plot([DM_B29X(1:Xmax) DM_B29X(Xmax)],[DM_B29Z(1:Xmax) 0],'color',[0.5 0.5 0.5],'LineWidth',taille_ligne)
z_DM = [z_DM mean(DM_B29Z(1:Xmax))];
x_DM = [x_DM DM_B29X(Xmax)];
n=n+1;
hold on

DM_B30X = [DM_B30.x];
DM_B30Z = [DM_B30.z];
Xmax = 33; % valeur de X avant premier crash
plot([DM_B30X(1:Xmax) DM_B30X(Xmax)],[DM_B30Z(1:Xmax) 0],'color',[0.5 0.5 0.5],'LineWidth',taille_ligne)
z_DM = [z_DM mean(DM_B30Z(1:Xmax))];
x_DM = [x_DM DM_B30X(Xmax)];
n=n+1;
hold on

DM_B31X = [DM_B31.x];
DM_B31Z = [DM_B31.z];
Xmax = 61; % valeur de X avant premier crash
plot([DM_B31X(1:Xmax) DM_B31X(Xmax)],[DM_B31Z(1:Xmax) 0],'color',[0.5 0.5 0.5],'LineWidth',taille_ligne)
z_DM = [z_DM mean(DM_B31Z(1:Xmax))];
x_DM = [x_DM DM_B31X(Xmax)];
n=n+1;
hold on

DM_B32X = [DM_B32.x];
DM_B32Z = [DM_B32.z];
Xmax = find(DM_B32Z==min(DM_B32Z)); % valeur de X avant premier crash
plot([DM_B32X(1:Xmax) DM_B32X(Xmax)],[DM_B32Z(1:Xmax) 0],'color',[0.5 0.5 0.5],'LineWidth',taille_ligne)
z_DM = [z_DM mean(DM_B32Z(1:Xmax))];
x_DM = [x_DM DM_B32X(Xmax)];
n=n+1;
hold on

DM_B33X = [DM_B33.x];
DM_B33Z = [DM_B33.z];
Xmax = find(DM_B33Z==min(DM_B33Z)); % valeur de X avant premier crash
plot([DM_B33X(1:Xmax) DM_B33X(Xmax)],[DM_B33Z(1:Xmax) 0],'color',[0.5 0.5 0.5],'LineWidth',taille_ligne)
z_DM = [z_DM mean(DM_B33Z(1:Xmax))];
x_DM = [x_DM DM_B33X(Xmax)];
n=n+1;
hold on

DM_B34X = [DM_B34.x];
DM_B34Z = [DM_B34.z];
Xmax = 52; % valeur de X avant premier crash
plot([DM_B34X(1:Xmax) DM_B34X(Xmax)],[DM_B34Z(1:Xmax) 0],'color',[0.5 0.5 0.5],'LineWidth',taille_ligne)
z_DM = [z_DM mean(DM_B34Z(1:Xmax))];
x_DM = [x_DM DM_B34X(Xmax)];
n=n+1;
hold on

DM_B35X = [DM_B35.x];
DM_B35Z = [DM_B35.z];
Xmax = 52; % valeur de X avant premier crash
plot([DM_B35X(1:Xmax) DM_B35X(Xmax)],[DM_B35Z(1:Xmax) 0],'color',[0.5 0.5 0.5],'LineWidth',taille_ligne)
z_DM = [z_DM mean(DM_B35Z(1:Xmax))];
x_DM = [x_DM DM_B35X(Xmax)];
n=n+1;
hold on

DM_B36X = [DM_B36.x];
DM_B36Z = [DM_B36.z];
Xmax = find(DM_B36Z==min(DM_B36Z)); % valeur de X avant premier crash
plot([DM_B36X(1:Xmax) DM_B36X(Xmax)],[DM_B36Z(1:Xmax) 0],'color',[0.5 0.5 0.5],'LineWidth',taille_ligne)
z_DM = [z_DM mean(DM_B36Z(1:Xmax))];
x_DM = [x_DM DM_B36X(Xmax)];
n=n+1;
hold on

DM_B37X = [DM_B37.x];
DM_B37Z = [DM_B37.z];
Xmax = 67; % valeur de X avant premier crash
plot([DM_B37X(1:Xmax) DM_B37X(Xmax)],[DM_B37Z(1:Xmax) 0],'color',[0.5 0.5 0.5],'LineWidth',taille_ligne)
z_DM = [z_DM mean(DM_B37Z(1:Xmax))];
x_DM = [x_DM DM_B37X(Xmax)];
n=n+1;
hold on


xticks([0 1.3]) % On programme l'échelle à afficher en x
yticks([-0.02 0 0.09 0.14 0.71]) % On programme l'échelle à afficher en y
xticklabels({'0','1.3'}) % On programme le texte à écrire sous l'échelle en x
yticklabels({'','0','0.09','0.14','0.71'}) % On programme le texte à écrire sous l'échelle en y
axis([0 1.3 -0.02 0.73]) % On donne la limite des axes
text(0.1,0.6,'(b) Double mirror condition') % On demande à écrire "(f)" en haut à gauche dans le graphique
str={n};
text(0.7,0.6,'n = ')
text(0.77,0.6,str)

% Dessin des motifs et miroirs dans le graphique
offsety = 7.1; % 100 mm

% 2 épaisseur

for p = 1 : 55
        m = (p-0.03)*0.04+0.01 ;
        hold on
       plot([m m-0.05 ],[0.73 0.71],'-b','LineWidth',1.3)
end
hold on 
plot([0 1.3], [0.71 0.71],'-b','LineWidth',1.3)
    
% 110 x 2 = 220 distance max

for p = 1 : 55
        m = (p-0.03)*0.04+0.01 ;
        hold on
       plot([m-0.2 m+0.1 ],[0 -0.2],'-b','LineWidth',1.3)
end
hold on 
plot([0 1.3], [0 0],'-b','LineWidth',1.3)

ylabel('Altitude (m)')
xlabel('Distance travelled (m)')
title('1st flight recorded in double mirror condition','FontWeight','normal')
box on

% 1er vol en demi miroir bas DMB
text(0.7,0.5,'z = 7.3 \pm 1.7 cm')
text(0.7,0.4,'x = 20.7 \pm 13.1 cm')

% 'z_DM'
% mean(z_DM*100)
% std(z_DM*100)
% 
% 'x_DM'
% mean(x_DM*100)
% std(x_DM*100)
% 
% print -f2 -dpng -r300 DM
%
%%%%%%%%%%%%%%%%%%%%%%%%%%%%%%%%%%%%%%%%%%%%%%%%%%%%%%%%%%%%%%%%%%%%%%%%%
%-------------------- CONTROLE MIROIR BAS ---------------------------------

% Trajectoires en condition contrôle (premier passage) avec les abeilles du groupe miroir bas : f6 (c)
figure(3)
subplot(2,1,1);

% Moyennes de chaques trajectoires (utiles pour faire les boîtes à moustaches)
z_CC1_MB = [];
K = 14/19; % facteur de calibration en Z
n=0;

%Tracé des trajectoires :
plot([CC1_B38.x]*130/(max([CC1_B38.x])*100),[CC1_B38.z]*K,'color',[0.5 0.5 0.5],'LineWidth',taille_ligne)
z_CC1_MB = [z_CC1_MB mean([CC1_B38.z]*K)];

n=n+1;
hold on

plot([CC1_B39.x]*130/(max([CC1_B39.x])*100),[CC1_B39.z]*K,'color',[0.5 0.5 0.5],'LineWidth',taille_ligne)
z_CC1_MB = [z_CC1_MB mean([CC1_B39.z]*K)];
n=n+1;
hold on

plot([CC1_B40.x]*130/(max([CC1_B40.x])*100),[CC1_B40.z]*K,'color',[0.5 0.5 0.5],'LineWidth',taille_ligne)
z_CC1_MB = [z_CC1_MB mean([CC1_B40.z]*K)];
n=n+1;
hold on

plot([CC1_B41.x]*130/(max([CC1_B41.x])*100),[CC1_B41.z]*K,'color',[0.5 0.5 0.5],'LineWidth',taille_ligne)
z_CC1_MB = [z_CC1_MB mean([CC1_B41.z]*K)];
n=n+1;
hold on

plot([CC1_B42.x]*130/(max([CC1_B42.x])*100),[CC1_B42.z]*K,'color',[0.5 0.5 0.5],'LineWidth',taille_ligne)
z_CC1_MB = [z_CC1_MB mean([CC1_B42.z]*K)];
n=n+1;
hold on

plot([CC1_B44.x]*130/(max([CC1_B44.x])*100),[CC1_B44.z]*K,'color',[0.5 0.5 0.5],'LineWidth',taille_ligne)
z_CC1_MB = [z_CC1_MB mean([CC1_B44.z]*K)];
n=n+1;
hold on

plot([CC1_B45.x]*130/(max([CC1_B45.x])*100),[CC1_B45.z]*K,'color',[0.5 0.5 0.5],'LineWidth',taille_ligne)
z_CC1_MB = [z_CC1_MB mean([CC1_B45.z]*K)];
n=n+1;
hold on

plot([CC1_B46.x]*130/(max([CC1_B46.x])*100),[CC1_B46.z]*K,'color',[0.5 0.5 0.5],'LineWidth',taille_ligne)
z_CC1_MB = [z_CC1_MB mean([CC1_B46.z]*K)];
n=n+1;
hold on

plot([CC1_B47.x]*130/(max([CC1_B47.x])*100),[CC1_B47.z]*K,'color',[0.5 0.5 0.5],'LineWidth',taille_ligne)
z_CC1_MB = [z_CC1_MB mean([CC1_B47.z]*K)];
n=n+1;
hold on

plot([CC1_B48.x]*130/(max([CC1_B48.x])*100),[CC1_B48.z]*K,'color',[0.5 0.5 0.5],'LineWidth',taille_ligne)
z_CC1_MB = [z_CC1_MB mean([CC1_B48.z]*K)];
n=n+1;
hold on

plot([CC1_B50.x]*130/(max([CC1_B50.x])*100),[CC1_B50.z]*K,'color',[0.5 0.5 0.5],'LineWidth',taille_ligne)
z_CC1_MB = [z_CC1_MB mean([CC1_B50.z]*K)];
n=n+1;
hold on

plot([CC1_B51.x]*130/(max([CC1_B51.x])*100),[CC1_B51.z]*K,'color',[0.5 0.5 0.5],'LineWidth',taille_ligne)
z_CC1_MB = [z_CC1_MB mean([CC1_B51.z]*K)];
n=n+1;
hold on

plot([CC_B53.x]*130/(max([CC_B53.x])*100),[CC_B53.z]*K,'color',[0.5 0.5 0.5],'LineWidth',taille_ligne)
z_CC1_MB = [z_CC1_MB mean([CC_B53.z]*K)];
n=n+1;
hold on

plot([CC_B55.x]*130/(max([CC_B55.x])*100),[CC_B55.z]*K,'color',[0.5 0.5 0.5],'LineWidth',taille_ligne)
z_CC1_MB = [z_CC1_MB mean([CC_B55.z]*K)];
n=n+1;
hold on

plot([CC_B56.x]*130/(max([CC_B56.x])*100),[CC_B56.z]*K,'color',[0.5 0.5 0.5],'LineWidth',taille_ligne)
z_CC1_MB = [z_CC1_MB mean([CC_B56.z]*K)];
n=n+1;

% Mise en forme du graphique :
ylabel('Altitude (m)')
title('16th flight recorded in control condition after training','FontWeight','normal')
box on
text(0.7,0.5,'z = 11.6 \pm 4.0 cm')

xticks([0 1.3])
yticks([-0.02 0 0.09 0.14 0.71]) 
xticklabels({'0','1.3'}) 
yticklabels({'','0','0.09','0.14','0.71'}) 
axis([0 1.3 -0.02 0.73])
text(0.1,0.6,'(a) Control condition')
str={n};
text(0.7,0.6,'n = ')
text(0.77,0.6,str)
% t=0.01*round(median(ToF_CC_MB)*100);
% text(1,0.55,strcat('ToF =',num2str(t),'  s'))


% Dessin des motifs et miroirs dans le graphique
offsety = 7.1; % 100 mm

% 2 épaisseur

for p = 1 : 22
        m = (p-0.01)*0.1-0.1 ;
        rectangle('Position',[m,offsety/10,3,0.2],'FaceColor','w','EdgeColor',[1 0 0]);   %
        rectangle('Position',[0.03+m,offsety/10,1,0.2],'FaceColor','r','EdgeColor',[1 0 0]); %
        rectangle('Position',[0.04+m,offsety/10,1,0.2],'FaceColor','w','EdgeColor',[1 0 0]); % 11 periode plafond
        rectangle('Position',[0.05+m,offsety/10,1,0.2],'FaceColor','r','EdgeColor',[1 0 0]); %
        rectangle('Position',[0.06+m,offsety/10,1,0.2],'FaceColor','w','EdgeColor',[1 0 0]); %
        rectangle('Position',[0.07+m,offsety/10,3,0.2],'FaceColor','r','EdgeColor',[1 0 0]); %
end

for p = 1 : 22
        m = (p-0.01)*0.1-0.1 ;
        rectangle('Position',[m,-0.2,3,0.2],'FaceColor','w','EdgeColor',[1 0 0]);   %
        rectangle('Position',[0.03+m,-0.2,1,0.2],'FaceColor','r','EdgeColor',[1 0 0]); %
        rectangle('Position',[0.04+m,-0.2,1,0.2],'FaceColor','w','EdgeColor',[1 0 0]); % 11 periode plafond
        rectangle('Position',[0.05+m,-0.2,1,0.2],'FaceColor','r','EdgeColor',[1 0 0]); %
        rectangle('Position',[0.06+m,-0.2,1,0.2],'FaceColor','w','EdgeColor',[1 0 0]); %
        rectangle('Position',[0.07+m,-0.2,3,0.2],'FaceColor','r','EdgeColor',[1 0 0]); %
end

'z_CC1_MB'
mean(z_CC1_MB)
std(z_CC1_MB)

%--------------------------- MIROIR BAS ------------------------------------
% Trajectoires en condition miroir bas : f7 (d)
subplot(2,1,2);

% Moyennes de chaques trajectoires (utiles pour faire les boîtes à % moustaches)
z_MB = [];
x_MB = [];

n=0;
%Tracé des trajectoires :

MB_B38X = [MB_B38.x];
MB_B38Z = [MB_B38.z]*K;
plot(MB_B38X(1:79),MB_B38Z(1:79),'color',[0.5 0.5 0.5],'LineWidth',taille_ligne)
z_MB = [z_MB mean(MB_B38Z(1:79))];
x_MB = [x_MB MB_B38X(79)];
n=n+1;
hold on

MB_B39X = [MB_B39.x];
MB_B39Z = [MB_B39.z]*K;
plot([MB_B39X(1:97) MB_B39X(97)],[MB_B39Z(1:97) 0],'color',[0.5 0.5 0.5],'LineWidth',taille_ligne)
z_MB = [z_MB mean(MB_B39Z(1:97))];
x_MB = [x_MB MB_B39X(97)];
n=n+1;
hold on

MB_B40X = [MB_B40.x];
MB_B40Z = [MB_B40.z]*K;
plot([MB_B40X(1:102) MB_B40X(102)],[MB_B40Z(1:102) 0],'color',[0.5 0.5 0.5],'LineWidth',taille_ligne)
z_MB = [z_MB mean(MB_B40Z(1:102))];
x_MB = [x_MB MB_B40X(102)];
n=n+1;
hold on

MB_B41X = [MB_B41.x];
MB_B41Z = [MB_B41.z]*K;
plot([MB_B41X(1:55) MB_B41X(55)],[MB_B41Z(1:55) 0],'color',[0.5 0.5 0.5],'LineWidth',taille_ligne)
z_MB = [z_MB mean(MB_B41Z(1:55))];
x_MB = [x_MB MB_B41X(55)];
n=n+1;
hold on

MB_B42X = [MB_B42.x];
MB_B42Z = [MB_B42.z]*K;
plot([MB_B42X(1:24) MB_B42X(24)],[MB_B42Z(1:24) 0],'color',[0.5 0.5 0.5],'LineWidth',taille_ligne)
z_MB = [z_MB mean(MB_B42Z(1:24))];
x_MB = [x_MB MB_B42X(24)];
n=n+1;
hold on

MB_B44X = [MB_B44.x];
MB_B44Z = [MB_B44.z]*K;
plot([MB_B44X(1:221) MB_B44X(221)],[MB_B44Z(1:221) 0],'color',[0.5 0.5 0.5],'LineWidth',taille_ligne)
z_MB = [z_MB mean(MB_B44Z(1:221))];
x_MB = [x_MB MB_B44X(221)];
n=n+1;
hold on

MB_B45X = [MB_B45.x];
MB_B45Z = [MB_B45.z]*K;
plot([MB_B45X(1:29) MB_B45X(29)],[MB_B45Z(1:29) 0],'color',[0.5 0.5 0.5],'LineWidth',taille_ligne)
z_MB = [z_MB mean(MB_B45Z(1:29))];
x_MB = [x_MB MB_B45X(29)];
n=n+1;
hold on

MB_B46X = [MB_B46.x];
MB_B46Z = [MB_B46.z]*K;
plot([MB_B46X(1:55) MB_B46X(55)],[MB_B46Z(1:55) 0],'color',[0.5 0.5 0.5],'LineWidth',taille_ligne)
z_MB = [z_MB mean(MB_B46Z(1:55))];
x_MB = [x_MB MB_B46X(55)];
n=n+1;
hold on


MB_B47X = [MB_B47.x];
MB_B47Z = [MB_B47.z]*K;
plot([MB_B47X(1:100) MB_B47X(100)],[MB_B47Z(1:100) 0],'color',[0.5 0.5 0.5],'LineWidth',taille_ligne)
z_MB = [z_MB mean(MB_B47Z(1:100))];
x_MB = [x_MB MB_B47X(100)];
n=n+1;
hold on

MB_B48X = [MB_B48.x];
MB_B48Z = [MB_B48.z]*K;
plot([MB_B48X(1:end) MB_B48X(end)],[MB_B48Z(1:end) 0],'color',[0.5 0.5 0.5],'LineWidth',taille_ligne)
z_MB = [z_MB mean(MB_B48Z(1:end))];
x_MB = [x_MB MB_B48X(end)];
n=n+1;
hold on

MB_B49X = [MB_B49.x];
MB_B49Z = [MB_B49.z]*K;
plot([MB_B49X(1:206) MB_B49X(206)],[MB_B49Z(1:206) 0],'color',[0.5 0.5 0.5],'LineWidth',taille_ligne)
z_MB = [z_MB mean(MB_B49Z(1:206))];
x_MB = [x_MB MB_B49X(206)];
n=n+1;
hold on

MB_B50X = [MB_B50.x];
MB_B50Z = [MB_B50.z]*K;
plot([MB_B50X(1:61) MB_B50X(61)],[MB_B50Z(1:61) 0],'color',[0.5 0.5 0.5],'LineWidth',taille_ligne)
z_MB = [z_MB mean(MB_B50Z(1:61))];
x_MB = [x_MB MB_B50X(61)];
n=n+1;
hold on

MB_B51X = [MB_B51.x];
MB_B51Z = [MB_B51.z]*K;
plot([MB_B51X(1:161) MB_B51X(161)],[MB_B51Z(1:161) 0],'color',[0.5 0.5 0.5],'LineWidth',taille_ligne)
z_MB = [z_MB mean(MB_B51Z(1:161))];
x_MB = [x_MB MB_B51X(161)];
n=n+1;
hold on

MB_B55X = [MB_B55.x];
MB_B55Z = [MB_B55.z]*K;
plot([MB_B55X(1:73) MB_B55X(73)],[MB_B55Z(1:73) 0],'color',[0.5 0.5 0.5],'LineWidth',taille_ligne)
z_MB = [z_MB mean(MB_B55Z(1:73))];
x_MB = [x_MB MB_B55X(73)];
n=n+1;
hold on

MB_B56X = [MB_B56.x];
MB_B56Z = [MB_B56.z]*K;
plot([MB_B56X(1:26) MB_B56X(26)],[MB_B56Z(1:26) 0],'color',[0.5 0.5 0.5],'LineWidth',taille_ligne)
z_MB = [z_MB mean(MB_B56Z(1:26))];
x_MB = [x_MB MB_B56X(26)];
n=n+1;


xticks([0 1.3])
yticks([-0.02 0 0.09 0.14 0.71])
xticklabels({'0','1.3'})
yticklabels({'','0','0.09','0.14','0.71'})
axis([0 1.3 -0.02 0.73]) 
text(0.1,0.6,'(b) Mirror on the floor condition') 
str={n};
text(0.7,0.6,'n = ')
text(0.77,0.6,str)

% Dessin des motifs et miroirs dans le graphique
offsety = 7.1; % 100 mm

% 2 épaisseur

for p = 1 : 22
        m = (p-0.01)*0.1-0.1 ;
        rectangle('Position',[m,offsety/10,3,0.2],'FaceColor','w','EdgeColor',[1 0 0]);   %
        rectangle('Position',[0.03+m,offsety/10,1,0.2],'FaceColor','r','EdgeColor',[1 0 0]); %
        rectangle('Position',[0.04+m,offsety/10,1,0.2],'FaceColor','w','EdgeColor',[1 0 0]); % 11 periode plafond
        rectangle('Position',[0.05+m,offsety/10,1,0.2],'FaceColor','r','EdgeColor',[1 0 0]); %
        rectangle('Position',[0.06+m,offsety/10,1,0.2],'FaceColor','w','EdgeColor',[1 0 0]); %
        rectangle('Position',[0.07+m,offsety/10,3,0.2],'FaceColor','r','EdgeColor',[1 0 0]); %
end
    
% 110 x 2 = 220 distance max

for p = 1 : 55
        m = (p-0.03)*0.04+0.01 ;
        hold on
       plot([m-0.2 m+0.1 ],[0 -0.2],'-b','LineWidth',1.3)
end
hold on 
plot([0 1.3], [0 0],'-b','LineWidth',1.3)

ylabel('Altitude (m)')
xlabel('Distance travelled (m)')
title('1st flight recorded in mirror on the floor condition','FontWeight','normal')
box on

% 1er vol en demi miroir bas DMB
text(0.7,0.5,'z = 8.4 \pm 2.1 cm')
text(0.7,0.4,'x = 33.1 \pm 22.5 cm')

print -f3 -dpng -r300 MB
```
