## Supplementary material for "An innovative optical context to make honeybees crash repeatedly": Data set of honeybees flights: script2_FigA.m

```
clear DMH_CCZ DMH_MHZ
%close(10)
RGB =rgb('LightCoral');
ymax = 0.71 ; % valeur maximale d'ordonnée sur le graphique
thickness = 0.02 ; % épaisseur graphisme environnement
pas = 0.08; % 7.5cm ou 7cm bien
taille = 1.3/pas; % 13
% Calibration de chacune des trajectoires
CCX01=[CC_B01.x]*128/163;
CCZ01=[CC_B01.z]*9/17;

CCX02=[CC_B02.x]*130/163;
CCZ02=[CC_B02.z]*9/14;

CCX03=[CC_B03.x]*130/163;
CCZ03=[CC_B03.z]*12/18;

CCX04=[CC_B04.x]*124/156;
CCZ04=[CC_B04.z]*9/16;

CCX05=[CC_B05.x]*130/163;
CCZ05=[CC_B05.z]*9/14;

CCX06=[CC_B06.x]*130/163;
CCZ06=[CC_B06.z]*9/11;

CCX07=[CC_B07.x]*130/164 ;
CCZ07=[CC_B07.z]*9/12 ;

CCX08=[CC_B08.x]*85/114;
CCZ08=[CC_B08.z]*9/12;

CCX09=[CC_B09.x]*130/164;
CCZ09=[CC_B09.z];

CCX10=[CC_B10.x]*128/164;
CCZ10=[CC_B10.z]*10/13;

CCX11=[CC_B11.x]*130/161;
CCZ11=[CC_B11.z]*9/11;

CCX12=[CC_B12.x]*130/164;
CCZ12=[CC_B12.z]*10/13;

CCX13=[CC_B13.x]*120/167;
CCZ13=[CC_B13.z]*9/14;

CCX14=[CC_B14.x]*66/102;
CCZ14=[CC_B14.z]*9/13;

CCX15=[CC_B15.x]*66/102;
CCZ15=[CC_B15.z]*9/13;

CCX16=[CC_B16.x]*130/174;
CCZ16=[CC_B16.z]*10/15;

CCX17=[CC_B17.x]*130/173;
CCZ17=[CC_B17.z]*10/14;

CCX18=[CC_B18.x]*130/173;
CCZ18=[CC_B18.z]*9/13;

CCX19=[CC_B19.x]*90/129;
CCZ19=[CC_B19.z]*12/17;

CCX20=[CC_B20.x]*130/173;
CCZ20=[CC_B20.z]*10/14;

CCX21=[CC_B21.x]*130/173;
CCZ21=[CC_B21.z]*9/13;

CCX22=[CC_B22.x]*130/97;
CCZ22=[CC_B22.z]*9/14;

CCX76=[CC_B76.x]*124/154;
CCZ76=[CC_B76.z]*9/23;

CCX77=[CC_B77.x]*124/154;
CCZ77=[CC_B77.z]*12/26;

CCX79=[CC_B79.x]*124/153;
CCZ79=[CC_B79.z]*9/24;

CCX80=[CC_B80.x]*124/155;
CCZ80=[CC_B80.z]*12/24;

CCX81=[CC_B81.x]*130/154;
CCZ81=[CC_B81.z]*9/23;

CCX82=[CC_B82.x]*124/155;
CCZ82=[CC_B82.z]*9/23;

X01 = [MH_B01.x]*130/164;
Z01 = [MH_B01.z]*10/17;

X02=[MH_B02.x]*130/164;
Z02=[MH_B02.z]*9/15;

X03=[MH_B03.x]*130/164;
Z03=[MH_B03.z]*12/19;

X04=[MH_B04.x]*130/164;
Z04=[MH_B04.z]*11/18;

X05=[MH_B05.x]*130/164;
Z05=[MH_B05.z]*10/16;

X06=[MH_B06.x]*130/164;
Z06=[MH_B06.z]*9/11;

X07=[MH_B07.x]*130/163;
Z07=[MH_B07.z]*9/12;

X08=[MH_B08.x]*36/60;
Z08=[MH_B08.z]*9/11;

X09=[MH_B09.x]*130/163;
Z09=[MH_B09.z]*12/14;

X10=[MH_B10.x]*130/163;
Z10=[MH_B10.z]*9/10;

X11=[MH_B11.x]*120/157;
Z11=[MH_B11.z]*9/11;

X12=[MH_B12.x]*52/80;
Z12=[MH_B12.z]*10/13;

X13=[MH_B13.x]*130/173;
Z13=[MH_B13.z]*12/16;

X14=[MH_B14.x]*130/173;
Z14=[MH_B14.z]*9/13;

X15=[MH_B15.x]*130/174;
Z15=[MH_B15.z]*11/15;
  
X16=[MH_B16.x]*130/173;
Z16=[MH_B16.z]*11/15;

X18=[MH_B18.x]*130/174;
Z18=[MH_B18.z]*11/16;

X19=[MH_B19.x]*6/16;
Z19=[MH_B19.z]*11/15;

X20=[MH_B20.x]*130/173;
Z20=[MH_B20.z]*11/16;

X21=[MH_B21.x]*130/173;
Z21=[MH_B21.z]*10/14;

X22=[MH_B22.x]*130/173;
Z22=[MH_B22.z]*10/14;

X76=[MH_B76.x]*130/155;
Z76=[MH_B76.z]*9/23;

X77=[MH_B77.x]*126/154;
Z77=[MH_B77.z]*11/23;

X79=[MH_B79.x]*130/155;
Z79=[MH_B79.z]*9/24;

X80=[MH_B80.x]*130/154;
Z80=[MH_B80.z]*8/21;

X81=[MH_B81.x]*130/154;
Z81=[MH_B81.z]*10/24;

X82=[MH_B82.x]*130/110;
Z82=[MH_B82.z]*3/16;
% DMH CCZ : Miroir Haut condition contrôle axe Z
DMH_CCZ(:,1)=binning(pas,CCX01,CCZ01);
DMH_CCZ(:,2)=binning(pas,CCX02,CCZ02);
DMH_CCZ(:,3)=binning(pas,CCX03,CCZ03);
DMH_CCZ(:,4)=binning(pas,CCX04,CCZ04);
DMH_CCZ(:,5)=binning(pas,CCX05,CCZ05);
DMH_CCZ(:,6)=binning(pas,CCX06,CCZ06);
DMH_CCZ(:,7)=binning(pas,CCX07,CCZ07);
DMH_CCZ(:,8)=binning(pas,CCX08,CCZ08);
DMH_CCZ(:,9)=binning(pas,CCX09,CCZ09);
DMH_CCZ(:,10)=binning(pas,CCX10,CCZ10);
DMH_CCZ(:,11)=binning(pas,CCX11,CCZ11);
DMH_CCZ(:,12)=binning(pas,CCX12,CCZ12);
DMH_CCZ(:,13)=binning(pas,CCX13,CCZ13);
DMH_CCZ(:,14)=binning(pas,CCX14,CCZ14);
DMH_CCZ(:,15)=binning(pas,CCX15,CCZ15);
DMH_CCZ(:,16)=binning(pas,CCX16,CCZ16);
%DMH_CCZ(:,17)=binning(pas,CCX17,CCZ17);
DMH_CCZ(:,17)=binning(pas,CCX18,CCZ18);
DMH_CCZ(:,18)=binning(pas,CCX19,CCZ19);
DMH_CCZ(:,19)=binning(pas,CCX20,CCZ20);
DMH_CCZ(:,20)=binning(pas,CCX21,CCZ21);
DMH_CCZ(:,21)=binning(pas,CCX22,CCZ22);
DMH_CCZ(:,22)=binning(pas,CCX76,CCZ76);
DMH_CCZ(:,23)=binning(pas,CCX77,CCZ77);
DMH_CCZ(:,24)=binning(pas,CCX79,CCZ79);
DMH_CCZ(:,25)=binning(pas,CCX80,CCZ80);
DMH_CCZ(:,26)=binning(pas,CCX81,CCZ81);
DMH_CCZ(:,27)=binning(pas,CCX82,CCZ82);

% DMH MHZ : Miroir Haut condition MH
DMH_MHZ(:,1)=binning(pas,X01,Z01);
DMH_MHZ(:,2)=binning(pas,X02,Z02);
DMH_MHZ(:,3)=binning(pas,X03,Z03);
DMH_MHZ(:,4)=binning(pas,X04,Z04);
DMH_MHZ(:,5)=binning(pas,X05,Z05);
DMH_MHZ(:,6)=binning(pas,X06,Z06);
DMH_MHZ(:,7)=binning(pas,X07,Z07);
DMH_MHZ(:,8)=binning(pas,X08,Z08);
DMH_MHZ(:,9)=binning(pas,X09,Z09);
DMH_MHZ(:,10)=binning(pas,X10,Z10);
DMH_MHZ(:,11)=binning(pas,X11,Z11);
DMH_MHZ(:,12)=binning(pas,X12,Z12);
DMH_MHZ(:,13)=binning(pas,X13,Z13);
DMH_MHZ(:,14)=binning(pas,X14,Z14);
DMH_MHZ(:,15)=binning(pas,X15,Z15);
DMH_MHZ(:,16)=binning(pas,X16,Z16);
%DMH_CCZ(:,17)=binning(pas,X17,Z17);
DMH_MHZ(:,17)=binning(pas,X18,Z18);
DMH_MHZ(:,18)=binning(pas,X19,Z19);
DMH_MHZ(:,19)=binning(pas,X20,Z20);
DMH_MHZ(:,20)=binning(pas,X21,Z21);
DMH_MHZ(:,21)=binning(pas,X22,Z22);
DMH_MHZ(:,22)=binning(pas,X76,Z76);
DMH_MHZ(:,23)=binning(pas,X77,Z77);
DMH_MHZ(:,24)=binning(pas,X79,Z79);
DMH_MHZ(:,25)=binning(pas,X80,Z80);
DMH_MHZ(:,26)=binning(pas,X81,Z81);
DMH_MHZ(:,27)=binning(pas,X82,Z82);
 

figure(10)
din = size(DMH_MHZ);
x_shade = pas*(0:(din(1)-1))+pas/2;
sp_hand = subplot(2,1,1)

shadedErrorBar(x_shade,DMH_CCZ',{@nanmedian,@madX},{'-ok','markerfacecolor','k','MarkerSize',4});
axis([0 2 -0.02 ymax+0.02])
xticks([0 1.3 2])%pour l'abscisse
xticklabels([])%pour l'abscisse
yticks([0 0.09 0.14 ymax 2*ymax])% positions des ticks sur l'ordonné
yticklabels({'0','0.09','0.14','0.71','1.42'}) % noms des ticks
title('16th flight recorded in control condition after training','FontWeight','normal')%titre du graph
%xlabel('Distance travelled (m)')
ylabel('Altitude (m)')
text(0.01,ymax-0.1,'(Aii) Control condition')
str='       n=15';
text(0.03,ymax-0.25,str)
hold on
%plot([0 0],[0.01 0.7*0.75],'color',[0 0 0],'LineStyle','--')
%plot([1.3 1.3],[0.01 0.7],'color',[0 0 0],'LineStyle','--')

% Dessin des motifs et miroirs dans le graphique
% 2 épaisseur
for p = 1 : 22
        m = (p-0.01)*0.1-0.1;
        rectangle('Position',[m,ymax,3,thickness],'FaceColor','w','EdgeColor',[1 0 0]);   %
        rectangle('Position',[0.03+m,ymax,1,thickness],'FaceColor','r','EdgeColor',[1 0 0]); %
        rectangle('Position',[0.04+m,ymax,1,thickness],'FaceColor','w','EdgeColor',[1 0 0]); % 11 periode plafond
        rectangle('Position',[0.05+m,ymax,1,thickness],'FaceColor','r','EdgeColor',[1 0 0]); %
        rectangle('Position',[0.06+m,ymax,1,thickness],'FaceColor','w','EdgeColor',[1 0 0]); %
        rectangle('Position',[0.07+m,ymax,3,thickness],'FaceColor','r','EdgeColor',[1 0 0]); %
end

for p = 1 : 22
        m = (p-0.01)*0.1-0.1;
        rectangle('Position',[m,-thickness,3,thickness],'FaceColor','w','EdgeColor',[1 0 0]);   %
        rectangle('Position',[0.03+m,-thickness,1,thickness],'FaceColor','r','EdgeColor',[1 0 0]); %
        rectangle('Position',[0.04+m,-thickness,1,thickness],'FaceColor','w','EdgeColor',[1 0 0]); % 11 periode plafond
        rectangle('Position',[0.05+m,-thickness,1,thickness],'FaceColor','r','EdgeColor',[1 0 0]); %
        rectangle('Position',[0.06+m,-thickness,1,thickness],'FaceColor','w','EdgeColor',[1 0 0]); %
        rectangle('Position',[0.07+m,-thickness,3,thickness],'FaceColor','r','EdgeColor',[1 0 0]); %
end
box on
axe = gca;
axe.FontSize=5;
pos1 = get(sp_hand, 'Position'); % gives the position of current sub-plot
    new_pos1 = [pos1(1) pos1(2)-0.04 pos1(3) pos1(4)/2];
    set(sp_hand, 'Position',new_pos1); % set new position of current sub - plot


subplot(2,1,2)% impoverished visual condition
% Dessin des motifs et miroirs dans le graphique
hold on
% 2 épaisseur
GrayWidth = 0.8; % largeur du motif du miroir
%Couleur_M = [0.5 0.5 0.5]; % gris
Couleur_M = [0 0 1]; % bleu

for p = 0 : 45
       m = 0.625/11*p+0; % 0 car on commance à 0
       plot([m m+0.625/11],[ymax+thickness ymax],'-','LineWidth',GrayWidth,'Color',Couleur_M)
end
hold on 

plot([0 2], [ymax ymax],'-','LineWidth',GrayWidth,'Color',Couleur_M)
plot([0 2], [ymax+thickness ymax+thickness],'-','LineWidth',GrayWidth,'Color',Couleur_M)
plot([0 0], [ymax+thickness ymax],'-','LineWidth',GrayWidth,'Color',Couleur_M)
plot([2 2], [ymax+thickness ymax],'-','LineWidth',GrayWidth,'Color',Couleur_M)

for p = 1 : 22
        m = (p-0.01)*0.1-0.1;
        rectangle('Position',[m,-thickness,3,thickness],'FaceColor','w','EdgeColor',[1 0 0]);   %
        rectangle('Position',[0.03+m,-thickness,1,thickness],'FaceColor','r','EdgeColor',[1 0 0]); %
        rectangle('Position',[0.04+m,-thickness,1,thickness],'FaceColor','w','EdgeColor',[1 0 0]); % 11 periode plafond
        rectangle('Position',[0.05+m,-thickness,1,thickness],'FaceColor','r','EdgeColor',[1 0 0]); %
        rectangle('Position',[0.06+m,-thickness,1,thickness],'FaceColor','w','EdgeColor',[1 0 0]); %
        rectangle('Position',[0.07+m,-thickness,3,thickness],'FaceColor','r','EdgeColor',[1 0 0]); %
end

couleur = [1 0 0 0.1];

for p = 1:22
        m = (p-0.01)*0.1-0.1;
        rectangle('Position',[m,2*ymax,3,thickness],'FaceColor','w','EdgeColor',couleur);   %
        rectangle('Position',[0.03+m,2*ymax,1,thickness],'FaceColor',couleur,'EdgeColor',couleur); %
        rectangle('Position',[0.04+m,2*ymax,1,thickness],'FaceColor','w','EdgeColor',couleur); % 11 periode plafond
        rectangle('Position',[0.05+m,2*ymax,1,thickness],'FaceColor',couleur,'EdgeColor',couleur); %
        rectangle('Position',[0.06+m,2*ymax,1,thickness],'FaceColor','w','EdgeColor',couleur); %
        rectangle('Position',[0.07+m,2*ymax,3,thickness],'FaceColor',couleur,'EdgeColor',couleur); %
        
        %rectangle('Position',[m,2*ymax,3,thickness],'FaceColor',couleur,'EdgeColor',couleur);
        %rectangle('Position',[0.03+m,2*ymax,1,thickness],'FaceColor','w','EdgeColor',couleur);
        %rectangle('Position',[0.06+m,2*ymax,1,thickness],'FaceColor',couleur,'EdgeColor',couleur);
        %rectangle('Position',[0.07+m,2*ymax,1,thickness],'FaceColor','w','EdgeColor',couleur);
        %rectangle('Position',[0.08+m,2*ymax,1,thickness],'FaceColor',couleur,'EdgeColor',couleur);
        %rectangle('Position',[0.09+m,2*ymax,3,thickness],'FaceColor','w','EdgeColor',couleur);
        %rectangle('Position',[0.10+m,2*ymax,3,thickness],'FaceColor',couleur,'EdgeColor',couleur);
end

shadedErrorBar(x_shade,DMH_MHZ',{@nanmedian,@madX},{'-ok','markerfacecolor','k','MarkerSize',4});
axis([0 2 -thickness 2*ymax+thickness])
%xticks([0 0.625 1.3]) % positions des ticks sur l'abscisse
xticks([0 1.3 2])%pour l'abscisse
yticks([0 0.09 0.14 ymax 2*ymax+thickness])% positions des ticks sur l'ordonné
yticklabels({'0','0.09','0.14','0.71','1.42'}) % noms des ticks
title('1st flight recorded ''in mirror on the ceiling'' condition','FontWeight','normal')%titre du graph
xlabel('Distance travelled (m)')
ylabel('Altitude (m)')
str='       n=15';
text(0.01,ymax-0.25,str)
text(0.01,ymax-0.1,'(Bii) Impoverished visual condition') 
axe = gca;
axe.FontSize=5;
box on

% Passage à l'impression en .png de la figure
print -f10 -dpng -r600 Mirroir_haut
% -f11 numéro de la figure
% -dpng format de la figure
% -r600 résolution
% ensuite c'est le nom
```
