## Supplementary material for "An innovative optical context to make honeybees crash repeatedly": Data set of honeybees flights: script2_FigB.m

```
clc
%close(7)
clear MB_CCZ MB_MBZ
RGB =rgb('LightCoral');
ymax = 0.71 ; % valeur maximale d'ordonnée sur le graphique
thickness = 0.02 ; % épaisseur graphisme environnement
pas = 0.08; % 7.5cm ou 7cm bien
taille = 1.3/pas; % 13
% MB CCZ : Miroir Bas condition contrôle axe Z 
MB_CCZ(:,1)=binning(pas,[CC1_B38.x]*130/(max([CC1_B38.x])*100),[CC1_B38.z]*K);
MB_CCZ(:,2)=binning(pas,[CC1_B39.x]*130/(max([CC1_B39.x])*100),[CC1_B39.z]*K);
MB_CCZ(:,3)=binning(pas,[CC1_B40.x]*130/(max([CC1_B40.x])*100),[CC1_B40.z]*K);
MB_CCZ(:,4)=binning(pas,[CC1_B41.x]*130/(max([CC1_B41.x])*100),[CC1_B41.z]*K);
MB_CCZ(:,5)=binning(pas,[CC1_B42.x]*130/(max([CC1_B42.x])*100),[CC1_B42.z]*K);
MB_CCZ(:,6)=binning(pas,[CC1_B44.x]*130/(max([CC1_B44.x])*100),[CC1_B44.z]*K);
MB_CCZ(:,7)=binning(pas,[CC1_B45.x]*130/(max([CC1_B45.x])*100),[CC1_B45.z]*K);
MB_CCZ(:,8)=binning(pas,[CC1_B46.x]*130/(max([CC1_B46.x])*100),[CC1_B46.z]*K);
MB_CCZ(:,9)=binning(pas,[CC1_B47.x]*130/(max([CC1_B47.x])*100),[CC1_B47.z]*K);
MB_CCZ(:,10)=binning(pas,[CC1_B48.x]*130/(max([CC1_B48.x])*100),[CC1_B48.z]*K);
MB_CCZ(:,11)=binning(pas,[CC1_B50.x]*130/(max([CC1_B50.x])*100),[CC1_B50.z]*K);
MB_CCZ(:,12)=binning(pas,[CC1_B51.x]*130/(max([CC1_B51.x])*100),[CC1_B51.z]*K);
MB_CCZ(:,13)=binning(pas,[CC_B53.x]*130/(max([CC_B53.x])*100),[CC_B53.z]*K);
MB_CCZ(:,14)=binning(pas,[CC_B55.x]*130/(max([CC_B55.x])*100),[CC_B55.z]*K);
MB_CCZ(:,15)=binning(pas,[CC_B56.x]*130/(max([CC_B56.x])*100),[CC_B56.z]*K);


%
MB_B38X = [MB_B38.x];% ancien vecteur X et Z
MB_B38Z = [MB_B38.z]*K;
X = MB_B38X(1:79);
Z = MB_B38Z(1:79);
clear  MB_B38X MB_B38Z 
MB_B38X = X; % nouveau vecteur X et Z
MB_B38Z  = Z;
clear X Z

MB_B39X = [MB_B39.x];
MB_B39Z = [MB_B39.z]*K;
X = [MB_B39X(1:97) MB_B39X(97)];
Z = [MB_B39Z(1:97) 0];
clear  MB_B39X MB_B39Z 
MB_B39X = X; % nouveau vecteur X et Z
MB_B39Z  = Z;
clear X Z

MB_B40X = [MB_B40.x];
MB_B40Z = [MB_B40.z]*K;
X = [MB_B40X(1:102) MB_B40X(102)];
Z = [MB_B40Z(1:102) 0];
clear  MB_B40X MB_B40Z 
MB_B40X = X; % nouveau vecteur X et Z
MB_B40Z  = Z;
clear X Z

MB_B41X = [MB_B41.x];
MB_B41Z = [MB_B41.z]*K;
X = [MB_B41X(1:55) MB_B41X(55)];
Z = [MB_B41Z(1:55) 0];
clear MB_B41X MB_B41Z
MB_B41X = X; % nouveau vecteur X et Z
MB_B41Z  = Z;
clear X Z

MB_B42X = [MB_B42.x];
MB_B42Z = [MB_B42.z]*K;
X = [MB_B42X(1:24) MB_B42X(24)];
Z = [MB_B42Z(1:24) 0];
clear MB_B42X MB_B42Z
MB_B42X = X; % nouveau vecteur X et Z
MB_B42Z  = Z;
clear X Z

MB_B44X = [MB_B44.x];
MB_B44Z = [MB_B44.z]*K;
X = [MB_B44X(1:221) MB_B44X(221)];
Z = [MB_B44Z(1:221) 0];
clear MB_B44X MB_B44Z
MB_B44X = X; % nouveau vecteur X et Z
MB_B44Z  = Z;
clear X Z

MB_B45X = [MB_B45.x];
MB_B45Z = [MB_B45.z]*K;
X = [MB_B45X(1:29) MB_B45X(29)];
Z = [MB_B45Z(1:29) 0];
clear MB_B45X MB_B45Z
MB_B45X = X; % nouveau vecteur X et Z
MB_B45Z  = Z;
clear X Z

MB_B46X = [MB_B46.x];
MB_B46Z = [MB_B46.z]*K;
X = [MB_B46X(1:55) MB_B46X(55)];
Z = [MB_B46Z(1:55) 0];
clear MB_B46X MB_B46Z
MB_B46X = X; % nouveau vecteur X et Z
MB_B46Z  = Z;
clear X Z

MB_B47X = [MB_B47.x];
MB_B47Z = [MB_B47.z]*K;
X = [MB_B47X(1:100) MB_B47X(100)];
Z = [MB_B47Z(1:100) 0];
clear MB_B47X MB_B47Z
MB_B47X = X; % nouveau vecteur X et Z
MB_B47Z  = Z;
clear X Z

MB_B48X = [MB_B48.x];
MB_B48Z = [MB_B48.z]*K;
X = [MB_B48X(1:end) MB_B48X(end)];
Z = [MB_B48Z(1:end) 0];
clear MB_B48X MB_B48Z
MB_B48X = X; % nouveau vecteur X et Z
MB_B48Z  = Z;
clear X Z

MB_B49X = [MB_B49.x];
MB_B49Z = [MB_B49.z]*K;
X = [MB_B49X(1:206) MB_B49X(206)];
Z = [MB_B49Z(1:206) 0];
clear MB_B49X MB_B49Z
MB_B49X = X; % nouveau vecteur X et Z
MB_B49Z  = Z;
clear X Z

MB_B50X = [MB_B50.x];
MB_B50Z = [MB_B50.z]*K;
X = [MB_B50X(1:61) MB_B50X(61)];
Z = [MB_B50Z(1:61) 0];
clear MB_B50X MB_B50Z
MB_B50X = X; % nouveau vecteur X et Z
MB_B50Z  = Z;
clear X Z

MB_B51X = [MB_B51.x];
MB_B51Z = [MB_B51.z]*K;
X = [MB_B51X(1:161) MB_B51X(161)];
Z = [MB_B51Z(1:161) 0];
clear MB_B51X MB_B51Z
MB_B51X = X; % nouveau vecteur X et Z
MB_B51Z  = Z;
clear X Z

MB_B55X = [MB_B55.x];
MB_B55Z = [MB_B55.z]*K;
X = [MB_B55X(1:73) MB_B55X(73)];
Z = [MB_B55Z(1:73) 0];
clear MB_B52X MB_B52Z
MB_B55X = X; % nouveau vecteur X et Z
MB_B55Z  = Z;
clear X Z

MB_B56X = [MB_B56.x];
MB_B56Z = [MB_B56.z]*K;
X = [MB_B56X(1:26) MB_B56X(26)];
Z = [MB_B56Z(1:26) 0];
clear MB_B56X MB_B56Z
MB_B56X = X; % nouveau vecteur X et Z
MB_B56Z  = Z;
clear X Z


% MB CCZ : Miroir Bas condition MB axe Z
MB_MBZ(:,1)=binning(pas,MB_B38X,MB_B38Z);
MB_MBZ(:,2)=binning(pas,MB_B39X,MB_B39Z);
MB_MBZ(:,3)=binning(pas,MB_B40X,MB_B40Z);
MB_MBZ(:,4)=binning(pas,MB_B41X,MB_B41Z);
MB_MBZ(:,5)=binning(pas,MB_B42X,MB_B42Z);
MB_MBZ(:,6)=binning(pas,MB_B44X,MB_B44Z);
MB_MBZ(:,7)=binning(pas,MB_B45X,MB_B45Z);
MB_MBZ(:,8)=binning(pas,MB_B46X,MB_B46Z);
MB_MBZ(:,9)=binning(pas,MB_B47X,MB_B47Z);
MB_MBZ(:,10)=binning(pas,MB_B48X,MB_B48Z);
MB_MBZ(:,11)=binning(pas,MB_B49X,MB_B49Z);
MB_MBZ(:,12)=binning(pas,MB_B50X,MB_B50Z);
MB_MBZ(:,13)=binning(pas,MB_B51X,MB_B51Z);
MB_MBZ(:,14)=binning(pas,MB_B55X,MB_B55Z);
MB_MBZ(:,15)=binning(pas,MB_B56X,MB_B56Z);


figure(7)
dim = size(MB_MBZ);
x_shade = pas*(0:(dim(1)-1))+pas/2;

sp_hand2 = subplot(2,1,1);
shadedErrorBar(x_shade,MB_CCZ',{@nanmedian,@madX},{'-ok','markerfacecolor','k','MarkerSize',4});
hold on
%plot([1.3 1.3],[0.01 0.7],'color','k','LineStyle','--')
axis([0 2 -0.02 ymax+0.02])
xticks([0 1.3 2])%pour l'abscisse
xticklabels([]) % noms des ticks
yticks([0 0.09 0.14 ymax])% positions des ticks sur l'ordonné
yticklabels({'0','0.09','0.14','0.71'}) % noms des ticks
title('16th flight recorded in control condition after training','FontWeight','normal')%titre du graph
%xlabel('Distance travelled (m)')
ylabel('Altitude (m)')
text(0.01,ymax-0.1,'(Aii) Control condition')
hold on
%plot([0 0],[0.01 0.7*0.75],'color',[0 0 0],'LineStyle','--')
%plot([1.3 1.3],[0.01 0.7],'color',[0 0 0],'LineStyle','--')
str='       n=15';
text(0.03,ymax-0.25,str)
% plot([0 0],[0.01 0.7*0.75],'color',[0 0 0],'LineStyle','--')
% plot([1.3 1.3],[0.01 0.7],'color',[0 0 0],'LineStyle','--')


% Dessin des motifs et miroirs dans le graphique

% 2 épaisseur
for p = 1 : 22
        m = (p-0.01)*0.1-0.1;
        rectangle('Position',[m,ymax,3,thickness],'FaceColor','w','EdgeColor',[1 0 0]);   %
        rectangle('Position',[0.03+m,ymax,1,thickness],'FaceColor','r','EdgeColor',[1 0 0]); %
        rectangle('Position',[0.04+m,ymax,1,thickness],'FaceColor','w','EdgeColor',[1 0 0]); % 11 periode plafond
        rectangle('Position',[0.05+m,ymax,1,thickness],'FaceColor','r','EdgeColor',[1 0 0]); %
        rectangle('Position',[0.06+m,ymax,1,thickness],'FaceColor','w','EdgeColor',[1 0 0]); %
        rectangle('Position',[0.07+m,ymax,3,thickness],'FaceColor','r','EdgeColor',[1 0 0]); %
end

for p = 1 : 22
        m = (p-0.01)*0.1-0.1;
        rectangle('Position',[m,-thickness,3,thickness],'FaceColor','w','EdgeColor',[1 0 0]);   %
        rectangle('Position',[0.03+m,-thickness,1,thickness],'FaceColor','r','EdgeColor',[1 0 0]); %
        rectangle('Position',[0.04+m,-thickness,1,thickness],'FaceColor','w','EdgeColor',[1 0 0]); % 11 periode plafond
        rectangle('Position',[0.05+m,-thickness,1,thickness],'FaceColor','r','EdgeColor',[1 0 0]); %
        rectangle('Position',[0.06+m,-thickness,1,thickness],'FaceColor','w','EdgeColor',[1 0 0]); %
        rectangle('Position',[0.07+m,-thickness,3,thickness],'FaceColor','r','EdgeColor',[1 0 0]); %
end
box on
axe = gca;
axe.FontSize=5;
pos1 = get(sp_hand2, 'Position'); % gives the position of current sub-plot
     new_pos1 = [pos1(1) pos1(2)-0.04 pos1(3) pos1(4)/2];
     set(sp_hand2, 'Position',new_pos1); % set new position of current sub - plot

%%%%%%%%%%% impoverished visual condition %%%%%%%%%%%     
subplot(2,1,2) 
hold on
% Dessin des motifs et miroirs dans le graphique

for p = 1 : 22
        m = (p-0.01)*0.1-0.1;
        rectangle('Position',[m,ymax,3,0.2],'FaceColor','w','EdgeColor',[1 0 0]);   %
        rectangle('Position',[0.03+m,ymax,1,0.2],'FaceColor','r','EdgeColor',[1 0 0]); %
        rectangle('Position',[0.04+m,ymax,1,0.2],'FaceColor','w','EdgeColor',[1 0 0]); % 11 periode plafond
        rectangle('Position',[0.05+m,ymax,1,0.2],'FaceColor','r','EdgeColor',[1 0 0]); %
        rectangle('Position',[0.06+m,ymax,1,0.2],'FaceColor','w','EdgeColor',[1 0 0]); %
        rectangle('Position',[0.07+m,ymax,3,0.2],'FaceColor','r','EdgeColor',[1 0 0]); %
end


GrayWidth = 0.8; % largeur du motif du miroir
%Couleur_M = [0.5 0.5 0.5]; % gris
Couleur_M = [0 0 1]; % bleu

for p = 0 : 45
       m = 0.625/11*p+0; % 0 car on commance à 0
       plot([m m+0.625/11],[0 -thickness],'-','LineWidth',GrayWidth,'Color',Couleur_M)
end

plot([0 2], [0 0],'-','LineWidth',GrayWidth,'Color','b')
plot([0 2],[-thickness -thickness],'color','b','LineWidth',GrayWidth)

% for p = 7 : 20
%         m = (p-0.01)*0.1-0.1;
%         rectangle('Position',[m,-thickness-ymax,3,thickness],'FaceColor','w','EdgeColor',RGB);   %
%         rectangle('Position',[0.03+m,-thickness-ymax,1,thickness],'FaceColor',RGB,'EdgeColor',RGB); %
%         rectangle('Position',[0.04+m,-thickness-ymax,1,thickness],'FaceColor','w','EdgeColor',RGB); % 11 periode plafond
%         rectangle('Position',[0.05+m,-thickness-ymax,1,thickness],'FaceColor',RGB,'EdgeColor',RGB); %
%         rectangle('Position',[0.06+m,-thickness-ymax,1,thickness],'FaceColor','w','EdgeColor',RGB); %
%         rectangle('Position',[0.07+m,-thickness-ymax,3,thickness],'FaceColor',RGB,'EdgeColor',RGB); %
% end

couleur = [1 0 0 0.1];

for p = 1:22
        m = (p-0.01)*0.1-0.1;
        rectangle('Position',[m,-ymax-thickness,3,thickness],'FaceColor','w','EdgeColor',couleur);   %
        rectangle('Position',[0.03+m,-ymax-thickness,1,thickness],'FaceColor',couleur,'EdgeColor',couleur); %
        rectangle('Position',[0.04+m,-ymax-thickness,1,thickness],'FaceColor','w','EdgeColor',couleur); % 11 periode plafond
        rectangle('Position',[0.05+m,-ymax-thickness,1,thickness],'FaceColor',couleur,'EdgeColor',couleur); %
        rectangle('Position',[0.06+m,-ymax-thickness,1,thickness],'FaceColor','w','EdgeColor',couleur); %
        rectangle('Position',[0.07+m,-ymax-thickness,3,thickness],'FaceColor',couleur,'EdgeColor',couleur); %
       
end
box on
        % plot([1.3 1.3],[-0.02-ymax ymax],'k','LineStyle','--')
        % plot([0 0],[-0.02-ymax ymax],'k','LineStyle','--')

shadedErrorBar(x_shade,MB_MBZ',{@nanmedian,@madX},{'-ok','markerfacecolor','k','MarkerSize',4});
axis([0 2 -0.02-ymax ymax+0.02])
xticks([0 1.3 2]) % positions des ticks sur l'abscisse
xticklabels({'0','1.3','2'}) % noms des ticks
yticks([-ymax-thickness 0 0.09 0.14 ymax])% positions des ticks sur l'ordonné
yticklabels({'-0.71','0','0.09','0.14','0.71'}) % noms des ticks
title('1st flight recorded in ''mirror on the floor'' condition','FontWeight','normal')%titre du graph
xlabel('Distance travelled (m)')
ylabel('Altitude (m)')
text(0.005,ymax-0.10,'(Bii) Impoverished visual condition') 
str='      n=15';
text(0.04,ymax-0.25,str)
axe = gca;
axe.FontSize=5;

% Passage à l'impression en .png de la figure
print -f7 -dpng -r600 Miroir_Bas
% -f9 numéro de la figure
% -dpng format de la figure
% -r600 résolution
% ensuite c'est le nom

return %point d'arrêt

%faire le test de normalité de Shapiro-Wilk pour la condition contrôle
for i=1:16
    vect_MB_CCZ(i)=mean(MB_CCZ(i,:),'omitnan')
end
%[H, pValue, W] = swtest(vect_MB_CCZ) %H=0 pValue=0.1474
% distribution normale des données

%faire le test de normalité de Shapiro-Wilk pour la condition MBZ
for i=1:16
    vect_MB_MBZ(i)=mean(MB_MBZ(i,:),'omitnan')
end
%[H, pValue, W] = swtest(vect_MB_MBZ)%H=0 Pvalue=0.6168 
% distribution normale des données

%les deux ont une distribution normale 
%vérification de l'homogénéité des var
 [h,pValue] = vartest2(vect_MB_CCZ,vect_MB_MBZ)
 %H=1 pValue=0.0117 les var ne sont pas homogènes
 
 %ttest pour var non homogène
 [h,pValue] = ttest2(vect_MB_CCZ,vect_MB_MBZ,'Vartype','unequal') 
 %il y a une différence significative h=1 pValue=0.0014
 
```
