## Supplementary material for "An innovative optical context to make honeybees crash repeatedly": Data set of honeybees flights: script2_FigC.m

```
clc
clear DM_CCZ DM_DMZ
RGB =rgb('LightCoral');
%close(8)
ymax = 0.71 ; % valeur maximale d'ordonnée sur le graphique
thickness = 0.02 ; % épaisseur graphisme environnement
pas = 0.08; % 7.5cm ou 7cm bien, 8cm le meilleur
taille = 1.3/pas; % 13
% DM CCZ :Double miroir condition contrôle axe Z (chercher la CC)
DM_CCZ(:,1)=binning(pas,[CC_B23.x]*130/max([CC_B23.x]*100),[CC_B23.z]);
DM_CCZ(:,2)=binning(pas,[CC_B24.x]*130/max([CC_B24.x]*100),[CC_B24.z]);
DM_CCZ(:,3)=binning(pas,[CC_B25.x]*130/max([CC_B25.x]*100),[CC_B25.z]);
DM_CCZ(:,4)=binning(pas,[CC_B26.x]*130/max([CC_B26.x]*100),[CC_B26.z]);
DM_CCZ(:,5)=binning(pas,[CC_B27.x]*130/max([CC_B27.x]*100),[CC_B27.z]);
DM_CCZ(:,6)=binning(pas,[CC_B28.x]*130/max([CC_B28.x]*100),[CC_B28.z]);
DM_CCZ(:,7)=binning(pas,[CC_B29.x]*130/max([CC_B29.x]*100),[CC_B29.z]);
DM_CCZ(:,8)=binning(pas,[CC_B30.x]*130/max([CC_B30.x]*100),[CC_B30.z]);
DM_CCZ(:,9)=binning(pas,[CC_B31.x]*130/max([CC_B31.x]*100),[CC_B31.z]);
DM_CCZ(:,10)=binning(pas,[CC_B32.x]*130/max([CC_B32.x]*100),[CC_B32.z]);
DM_CCZ(:,11)=binning(pas,[CC_B33.x]*130/max([CC_B33.x]*100),[CC_B33.z]);
DM_CCZ(:,12)=binning(pas,[CC_B34.x]*130/max([CC_B34.x]*100),[CC_B34.z]);
DM_CCZ(:,13)=binning(pas,[CC_B35.x]*130/max([CC_B35.x]*100),[CC_B35.z]);
DM_CCZ(:,14)=binning(pas,[CC_B37.x]*130/max([CC_B37.x]*100),[CC_B37.z]);
DM_CCZ(:,15)=binning(pas,[CC_B54.x]*130/max([CC_B54.x]*100),[CC_B54.z]);

% 
Xmax = 125; % valeur de X avant premier crash
DM_B23X = [DM_B23.x]; % ancien vecteur X et Z
DM_B23Z = [DM_B23.z];
X = [DM_B23X(1:Xmax) DM_B23X(Xmax)];
Z = [DM_B23Z(1:Xmax) 0];
clear DM_B23X DM_B23Z
DM_B23X = X; % nouveau vecteur X et Z
DM_B23Z = Z;
clear X Z

Xmax = 90; % valeur de X avant premier crash 
DM_B24X = [DM_B24.x];% ancien vecteur X et Z
DM_B24Z = [DM_B24.z];
X = [DM_B24X(1:Xmax) DM_B24X(Xmax)];
Z = [DM_B24Z(1:Xmax) 0];
clear DM_B24X DM_B24Z
DM_B24X= X; % nouveau vecteur X et Z
DM_B24Z = Z;
clear X Z

Xmax = 80; % valeur de X avant premier crash (20)
DM_B25X = [DM_B25.x];
DM_B25Z = [DM_B25.z];
X = [DM_B25X(1:Xmax) DM_B25X(Xmax)];
Z = [DM_B25Z(1:Xmax) 0];
clear DM_B25X DM_B25Z
DM_B25X= X; % nouveau vecteur X et Z
DM_B25Z = Z;
clear X Z


Xmax = 28; % valeur de X avant premier crash
DM_B26X = [DM_B26.x];% ancien vecteur X et Z
DM_B26Z = [DM_B26.z];
X =[DM_B26X(1:Xmax) DM_B26X(Xmax)];
Z = [DM_B26Z(1:Xmax) 0];
clear DM_B26X DM_B26Z 
DM_B26X = X; % nouveau vecteur X et Z
DM_B26Z = Z;
clear X Z

Xmax = find(DM_B27Z==min(DM_B27Z)); % valeur de X avant premier crash
DM_B27X = [DM_B27.x];% ancien vecteur X et Z
DM_B27Z = [DM_B27.z];
X = [DM_B27X(1:Xmax) DM_B27X(Xmax)];
Z = [DM_B27Z(1:Xmax) 0];
clear DM_B27X DM_B27Z 
DM_B27X = X; % nouveau vecteur X et Z
DM_B27Z = Z;
clear X Z


DM_B28X = [DM_B28.x];% ancien vecteur X et Z
DM_B28Z = [DM_B28.z];
Xmax = find(DM_B28Z==min(DM_B28Z)); % valeur de X avant premier crash
X = [DM_B28X(1:Xmax) DM_B28X(Xmax)];
Z = [DM_B28Z(1:Xmax) 0];
clear DM_B28X DM_B28Z 
DM_B28X = X; % nouveau vecteur X et Z
DM_B28Z = Z;
clear X Z

Xmax = find(DM_B29Z==min(DM_B29Z)); % valeur de X avant premier crash
DM_B29X = [DM_B29.x];% ancien vecteur X et Z
DM_B29Z = [DM_B29.z];
X = [DM_B29X(1:Xmax) DM_B29X(Xmax)];
Z = [DM_B29Z(1:Xmax) 0];
clear DM_B29X DM_B29Z 
DM_B29X = X; % nouveau vecteur X et Z
DM_B29Z = Z;
clear X Z


Xmax = 33; % valeur de X avant premier crash
DM_B30X = [DM_B30.x];% ancien vecteur X et Z
DM_B30Z = [DM_B30.z];
X = [DM_B30X(1:Xmax) DM_B30X(Xmax)];
Z = [DM_B30Z(1:Xmax) 0];
clear DM_B30X DM_B30Z 
DM_B30X = X; % nouveau vecteur X et Z
DM_B30Z = Z;
clear X Z

Xmax = 61; % valeur de X avant premier crash
DM_B31X = [DM_B31.x];
DM_B31Z = [DM_B31.z];
X = [DM_B31X(1:Xmax) DM_B31X(Xmax)];
Z = [DM_B31Z(1:Xmax) 0];
clear DM_B31X DM_B31Z 
DM_B31X = X; % nouveau vecteur X et Z
DM_B31Z = Z;
clear X Z

DM_B32X = [DM_B32.x];
DM_B32Z = [DM_B32.z];
Xmax = find(DM_B32Z==min(DM_B32Z)); % valeur de X avant premier crash
X = [DM_B32X(1:Xmax) DM_B32X(Xmax)];
Z = [DM_B32Z(1:Xmax) 0];
clear DM_B32X DM_B32Z 
DM_B32X = X; % nouveau vecteur X et Z
DM_B32Z = Z;
clear X Z


DM_B33X = [DM_B33.x];
DM_B33Z = [DM_B33.z];
Xmax = find(DM_B33Z==min(DM_B33Z)); % valeur de X avant premier crash
X = [DM_B33X(1:Xmax) DM_B33X(Xmax)];
Z = [DM_B33Z(1:Xmax) 0];
clear DM_B33X DM_B33Z 
DM_B33X = X; % nouveau vecteur X et Z
DM_B33Z = Z;
clear X Z

Xmax = 52; % valeur de X avant premier crash
DM_B34X = [DM_B34.x];
DM_B34Z = [DM_B34.z];
X = [DM_B34X(1:Xmax) DM_B34X(Xmax)];
Z = [DM_B34Z(1:Xmax) 0];
clear DM_B34X DM_B34Z 
DM_B34X = X; % nouveau vecteur X et Z
DM_B34Z = Z;
clear X Z

Xmax = 52; % valeur de X avant premier crash
DM_B35X = [DM_B35.x];
DM_B35Z = [DM_B35.z];
X = [DM_B35X(1:Xmax) DM_B35X(Xmax)];
Z = [DM_B35Z(1:Xmax) 0];
clear DM_B35X DM_B35Z 
DM_B35X = X; % nouveau vecteur X et Z
DM_B35Z = Z;
clear X Z

DM_B36X = [DM_B36.x];
DM_B36Z = [DM_B36.z];
Xmax = find(DM_B36Z==min(DM_B36Z)); % valeur de X avant premier crash
X = [DM_B36X(1:Xmax) DM_B36X(Xmax)];
Z = [DM_B36Z(1:Xmax) 0];
clear DM_B36X DM_B36Z 
DM_B36X = X; % nouveau vecteur X et Z
DM_B36Z = Z;
clear X Z

Xmax = 67; % valeur de X avant premier crash
DM_B37X = [DM_B37.x];
DM_B37Z = [DM_B37.z];
X = [DM_B37X(1:Xmax) DM_B37X(Xmax)];
Z = [DM_B37Z(1:Xmax) 0];
clear DM_B37X DM_B37Z 
DM_B37X = X; % nouveau vecteur X et Z
DM_B37Z = Z;
clear X Z

% DM CCZ : Double Miroir condition Double Miroir axe Z
DM_DMZ(:,1)=binning(pas,DM_B23X,DM_B23Z);
DM_DMZ(:,2)=binning(pas,DM_B24X,DM_B24Z);
DM_DMZ(:,3)=binning(pas,DM_B25X,DM_B25Z);
DM_DMZ(:,4)=binning(pas,DM_B26X,DM_B26Z);
DM_DMZ(:,5)=binning(pas,DM_B27X,DM_B27Z);
DM_DMZ(:,6)=binning(pas,DM_B28X,DM_B28Z);
DM_DMZ(:,7)=binning(pas,DM_B29X,DM_B29Z);
DM_DMZ(:,8)=binning(pas,DM_B30X,DM_B30Z);
DM_DMZ(:,9)=binning(pas,DM_B31X,DM_B31Z);
DM_DMZ(:,10)=binning(pas,DM_B32X,DM_B32Z);
DM_DMZ(:,11)=binning(pas,DM_B33X,DM_B33Z);
DM_DMZ(:,12)=binning(pas,DM_B34X,DM_B34Z);
DM_DMZ(:,13)=binning(pas,DM_B35X,DM_B35Z);
DM_DMZ(:,14)=binning(pas,DM_B36X,DM_B36Z);
DM_DMZ(:,15)=binning(pas,DM_B37X,DM_B37Z);


%figure(5)
%subplot(2,1,1)
%boxplot(DM_CCZ')

%subplot(2,1,2)
%boxplot(DM_DMZ')
%boxplot(DM_DMZ','PlotStyle','compact')

% MAD : mean absolute deviation = un paramètre de dispersion pour
% distribution anormale

figure(8)
dim = size(DM_DMZ);
x_shade = pas*(0:(dim(1)-1))+pas/2;

sp_hand3 = subplot(2,1,1);
shadedErrorBar(x_shade,DM_CCZ',{@nanmedian,@madX},{'-ok','markerfacecolor','k','MarkerSize',4});
hold on
%plot([1.3 1.3],[0.01 0.7],'color','k','LineStyle','--')
axis([0 2 -0.02 ymax+0.02])
xticks([0 1.3 2])%pour l'abscisse
xticklabels({'','',''}) % noms des ticks
yticks([0 0.09 0.14 ymax])% positions des ticks sur l'ordonné
yticklabels({'0','0.09','0.14','0.71'}) % noms des ticks
title('16th flight recorded in control condition after training','FontWeight','normal')%titre du graph
%xlabel('Distance travelled (m)')
ylabel('Altitude (m)')
text(0.01,ymax-0.1,'(Aii) Control condition') 
str='        n=15';
text(0.01,ymax-0.25,str)

% Dessin des motifs et miroirs dans le graphique

% 2 épaisseur
for p = 1 : 22
        m = (p-0.01)*0.1-0.1 ;
        rectangle('Position',[m,ymax,3,thickness],'FaceColor','w','EdgeColor',[1 0 0]);   %
        rectangle('Position',[0.03+m,ymax,1,thickness],'FaceColor','r','EdgeColor',[1 0 0]); %
        rectangle('Position',[0.04+m,ymax,1,thickness],'FaceColor','w','EdgeColor',[1 0 0]); % 11 periode plafond
        rectangle('Position',[0.05+m,ymax,1,thickness],'FaceColor','r','EdgeColor',[1 0 0]); %
        rectangle('Position',[0.06+m,ymax,1,thickness],'FaceColor','w','EdgeColor',[1 0 0]); %
        rectangle('Position',[0.07+m,ymax,3,thickness],'FaceColor','r','EdgeColor',[1 0 0]); %
end

for p = 1 : 22
        m = (p-0.01)*0.1-0.1 ;
        rectangle('Position',[m,-thickness,3,thickness],'FaceColor','w','EdgeColor',[1 0 0]);   %
        rectangle('Position',[0.03+m,-thickness,1,thickness],'FaceColor','r','EdgeColor',[1 0 0]); %
        rectangle('Position',[0.04+m,-thickness,1,thickness],'FaceColor','w','EdgeColor',[1 0 0]); % 11 periode plafond
        rectangle('Position',[0.05+m,-thickness,1,thickness],'FaceColor','r','EdgeColor',[1 0 0]); %
        rectangle('Position',[0.06+m,-thickness,1,thickness],'FaceColor','w','EdgeColor',[1 0 0]); %
        rectangle('Position',[0.07+m,-thickness,3,thickness],'FaceColor','r','EdgeColor',[1 0 0]); %
end
box on
axe = gca;
axe.FontSize=5;
pos1 = get(sp_hand3, 'Position'); % gives the position of current sub-plot
     new_pos1 = [pos1(1) pos1(2)-0.04 pos1(3) pos1(4)/2];
     set(sp_hand3, 'Position',new_pos1); % set new position of current sub - plot

%%%%%%%%%%% impoverished visual condition %%%%%%%%%%%
sp_hand4 = subplot(2,1,2);
hold on
% Dessin des motifs et miroirs dans le graphique

GrayWidth = 0.8; % largeur du motif du miroir

for p = 1 : 85
       m = (p-0.03)*0.04+0.01-0.5 ;
       plot([m-0.2 m+0.1],[0.91 0.71],'-','LineWidth',GrayWidth,'Color','b')
       plot([m-0.2 m+0.1],[0 -0.2],'-','LineWidth',GrayWidth,'Color','b')
end
plot([0 2], [0.71 0.71],'-','LineWidth',GrayWidth,'Color','b')
plot([0 2], [0.73 0.73],'-','LineWidth',GrayWidth,'Color','b')
plot([0 2], [0 0],'-','LineWidth',GrayWidth,'Color','b')
plot([0 2],[-thickness -thickness],'color','b','LineWidth',GrayWidth)

shadedErrorBar(x_shade,DM_DMZ',{@nanmedian,@madX},{'-ok','markerfacecolor','k','MarkerSize',4});
axis([0 2 -0.02 ymax+0.02])
%xticks([0 0.625 1.3]) % positions des ticks sur l'abscisse
xticks([0 1.3 2])%pour l'abscisse
yticks([0 0.09 0.14 ymax])% positions des ticks sur l'ordonné
yticklabels({'0','0.09','0.14','0.71'}) % noms des ticks
title('1st flight recorded ''in double mirror'' condition','FontWeight','normal')%titre du graph
xlabel('Distance travelled (m)')
ylabel('Altitude (m)')
text(0.01,ymax-0.1,'(Bii) Impoverished visual condition') 
str='        n=15';
text(0.01,ymax-0.25,str)
axe = gca;
axe.FontSize=5;
box on

pos2 = get(sp_hand4, 'Position'); % gives the position of current sub-plot
     new_pos2 = [pos2(1) pos2(2)+0.18 pos2(3) pos2(4)/2];
     set(sp_hand4, 'Position',new_pos2); % set new position of current sub - plot

% Passage à l'impression en .png de la figure
print -f8 -dpng -r600 Double_Miroir
% -f8 numéro de la figure
% -dpng format de la figure
% -r600 résolution
% ensuite c'est le nom
```
