## Supplementary material for "An innovative optical context to make honeybees crash repeatedly": Data set of honeybees flights: script2_FigD.m

```
clc
clear DMB_CCZ DMB_MBZ
RGB =rgb('LightCoral');
%close(9)
ymax = 0.71 ; % valeur maximale d'ordonnée sur le graphique
thickness = 0.02 ; % épaisseur graphisme environnement
pas = 0.08; % 7.5cm ou 7cm bien, 8cm meilleur
taille = 1.3/pas; % 13
% DMB CCZ : Demi-Miroir Bas condition contrôle axe Z
DMB_CCZ(:,1)=binning(pas,CCX60,CCZ60);
DMB_CCZ(:,2)=binning(pas,CCX61,CCZ61);
DMB_CCZ(:,3)=binning(pas,CCX62,CCZ62);
DMB_CCZ(:,4)=binning(pas,CCX63,CCZ63);
DMB_CCZ(:,5)=binning(pas,CCX64,CCZ64);
DMB_CCZ(:,6)=binning(pas,CCX65,CCZ65);
DMB_CCZ(:,7)=binning(pas,CCX67,CCZ67);
DMB_CCZ(:,8)=binning(pas,CCX68,CCZ68);
DMB_CCZ(:,9)=binning(pas,CCX69,CCZ69);
DMB_CCZ(:,10)=binning(pas,CCX72,CCZ72);
DMB_CCZ(:,11)=binning(pas,CCX73,CCZ73);
DMB_CCZ(:,12)=binning(pas,CCX74,CCZ74);
DMB_CCZ(:,13)=binning(pas,CCX75,CCZ75);
DMB_CCZ(:,14)=binning(pas,CCX78,CCZ78);

Z60 = [DMB_B60.z];
X60 = [DMB_B60.x]*130/(max([DMB_B66.x]*100));
X = X60(1:182);
Z = [Z60(1:181) 0];
clear X60 Z60
X60 = X; % nouveau vecteur X et Z
Z60 = Z;
clear X Z

Z61 = [DMB_B61.z];
X61 = [DMB_B61.x]*130/(max([DMB_B61.x]*100));
X = X61(1:163);
Z = [Z61(1:162) 0];
clear X61 Z61
X61 = X; 
Z61 = Z;
clear X Z

Z62 = [DMB_B62.z];
X62 = [DMB_B62.x];
X = X62(1:245);
Z = [Z62(1:244) 0];
clear X62 Z62
X62 = X;
Z62 = Z;
clear X Z 

Z63 = [DMB_B63.z];
X63 = [DMB_B63.x];
X = X63(1:186);
Z = [Z63(1:185) 0];
clear X63 Z63
X63 = X;
Z63 = Z;
clear X Z 

Z65 = [DMB_B65.z];
X65 = [DMB_B65.x];
X = X65(1:110);
Z = [Z65(1:109) 0];
clear X65 Z65
X65 = X; 
Z65 = Z;
clear X Z 

Z66 = [DMB_B66.z];
X66 = [DMB_B66.x]*130/(max([DMB_B66.x]*100));
X = X66(1:162); 
Z = [Z66(1:161) 0];
clear X66 Z66
X66 = X;
Z66 = Z;
clear X Z 

Z67 = [DMB_B67.z];
X67 = [DMB_B67.x]*130/(max([DMB_B67.x]*100));
X = X67(1:97);
Z = [Z67(1:96) 0];
clear X67 Z67
X67 = X;
Z67 = Z;
clear X Z

Z68 = [DMB_B68.z];
X68 = [DMB_B68.x];
X = X68(1:310);
Z = [Z68(1:309) 0];
clear X68 Z68
X68 = X;
Z68 = Z;
clear X Z

Z69 = [DMB_B69.z];
X69 = [DMB_B69.x]*130/(max([DMB_B69.x]*100));
X = X69(1:205);
Z = [Z69(1:204) 0];
clear X69 Z69
X69 = X;
Z69 = Z;
clear X Z

Z70 = [DMB_B70.z];
X70 = [DMB_B70.x];
X = X70(1:end);
Z = Z70(1:end);
clear X70 Z70
X70 = X;
Z70 = Z;
clear X Z

Z72 = [DMB_B72.z];
X72 = [DMB_B72.x]*130/(max([DMB_B72.x]*100));
X = X72(1:176);
Z = [Z72(1:175) 0];
clear X72 Z72
X72 = X;
Z72 = Z;
clear X Z

Z73 = [DMB_B73.z];
X73 = [DMB_B73.x]*130/(max([DMB_B72.x]*100));
X = X73(1:132);
Z = [Z73(1:131) 0];
clear X73 Z73
X73 = X;
Z73 = Z;
clear X Z

Z74 = [DMB_B74.z];
X74 = [DMB_B74.x]*130/(max([DMB_B74.x]*100));
X = X74(1:153);
Z = [Z74(1:152) 0];
clear X74 Z74
X74 = X;
Z74 = Z;
clear X Z

Z75 = [DMB_B75.z];
X75 = [DMB_B75.x];
X = X75(1:155);
Z = [Z75(1:154) 0];
clear X75 Z75
X75 = X;
Z75 = Z;
clear X Z

Z78 = [DMB_B78.z];
X78 = [DMB_B78.x];
X = X78(1:109);
Z = [Z78(1:108) 0];
clear X78 Z78
X78 = X;
Z78 = Z;
clear X Z


% DMB CCZ : Demi-Miroir Bas condition DMB axe Z
DMB_MBZ(:,1)=binning(pas,X60,Z60);
DMB_MBZ(:,2)=binning(pas,X61,Z61);
DMB_MBZ(:,3)=binning(pas,X62,Z62);
DMB_MBZ(:,4)=binning(pas,X63,Z63);
DMB_MBZ(:,6)=binning(pas,X65,Z65);
DMB_MBZ(:,6)=binning(pas,X66,Z66);
DMB_MBZ(:,7)=binning(pas,X67,Z67);
DMB_MBZ(:,8)=binning(pas,X68,Z68);
DMB_MBZ(:,9)=binning(pas,X69,Z69);
DMB_MBZ(:,9)=binning(pas,X70,Z70);
DMB_MBZ(:,10)=binning(pas,X72,Z72);
DMB_MBZ(:,11)=binning(pas,X73,Z73);
DMB_MBZ(:,12)=binning(pas,X74,Z74);
DMB_MBZ(:,13)=binning(pas,X75,Z75);
DMB_MBZ(:,14)=binning(pas,X78,Z78);


figure(9)
dimm = size(DMB_MBZ);
x_shade = pas*(0:(dimm(1)-1))+pas/2;

sp_hand1 = subplot(2,1,1)
shadedErrorBar(x_shade,DMB_CCZ',{@nanmedian,@madX},{'-ok','markerfacecolor','k','MarkerSize',4});
axis([-0.35 1.65 -0.02 ymax+0.02])
xticks([-0.35 0 0.625 1.3 1.65])%pour l'abscisse
xticklabels([0 0.35 1 1.65 2])%pour l'abscisse
yticks([0 0.09 0.14 ymax])% positions des ticks sur l'ordonné
yticklabels({'0','0.09','0.14','0.71'}) % noms des ticks
title('16th flight recorded in control condition after training','FontWeight','normal')%titre du graph
xlabel('Distance travelled (m)')
ylabel('Altitude (m)')
text(0.01-0.35,ymax-0.1,'(Aii) Control condition')
hold on
% traits verticaux
%plot([0.625 0.625],[0.01 0.7],'color',[0 0 0],'LineStyle','--')
%plot([0 0],[0.01 0.7*0.75],'color',[0 0 0],'LineStyle','--')
%plot([1.3 1.3],[0.01 0.7],'color',[0 0 0],'LineStyle','--')
str='      n=14';
text(0.03-0.35,ymax-0.25,str)

% Dessin des motifs et miroirs dans le graphique

for p = 1 : 22
        m = (p-0.01)*0.1-0.1-0.35 ; %-0.35 translation pour faire le zéro
        rectangle('Position',[m,ymax,3,thickness],'FaceColor','w','EdgeColor',[1 0 0]);   %
        rectangle('Position',[0.03+m,ymax,1,thickness],'FaceColor','r','EdgeColor',[1 0 0]); %
        rectangle('Position',[0.04+m,ymax,1,thickness],'FaceColor','w','EdgeColor',[1 0 0]); % 11 periode plafond
        rectangle('Position',[0.05+m,ymax,1,thickness],'FaceColor','r','EdgeColor',[1 0 0]); %
        rectangle('Position',[0.06+m,ymax,1,thickness],'FaceColor','w','EdgeColor',[1 0 0]); %
        rectangle('Position',[0.07+m,ymax,3,thickness],'FaceColor','r','EdgeColor',[1 0 0]); %
end

%plot([0 1.3], [ymax ymax],'-r','LineWidth',1.3)
%plot([0 1.3], [0 0],'-r','LineWidth',1.3)

for p = 1 : 22
        m = (p-0.01)*0.1-0.1-0.35 ; %-0.35 translation pour faire le zéro
        rectangle('Position',[m,-thickness,3,thickness],'FaceColor','w','EdgeColor',[1 0 0]);   %
        rectangle('Position',[0.03+m,-thickness,1,thickness],'FaceColor','r','EdgeColor',[1 0 0]); %
        rectangle('Position',[0.04+m,-thickness,1,thickness],'FaceColor','w','EdgeColor',[1 0 0]); % 11 periode plafond
        rectangle('Position',[0.05+m,-thickness,1,thickness],'FaceColor','r','EdgeColor',[1 0 0]); %
        rectangle('Position',[0.06+m,-thickness,1,thickness],'FaceColor','w','EdgeColor',[1 0 0]); %
        rectangle('Position',[0.07+m,-thickness,3,thickness],'FaceColor','r','EdgeColor',[1 0 0]); %
end
box on
axe = gca;
axe.FontSize=5;

pos1 = get(sp_hand1, 'Position'); % gives the position of current sub-plot
    new_pos1 = [pos1(1) pos1(2)-0.04 pos1(3) pos1(4)/2];
    set(sp_hand1, 'Position',new_pos1); % set new position of current sub - plot

subplot(2,1,2) % impoverished visual condition
% Dessin des motifs et miroirs dans le graphique
hold on 

for p = 1 : 22
        m = (p-0.01)*0.1-0.1-0.35 ; %-0.35 translation pour faire le zéro
        rectangle('Position',[m,ymax,3,thickness],'FaceColor','w','EdgeColor',[1 0 0]);   %
        rectangle('Position',[0.03+m,ymax,1,thickness],'FaceColor','r','EdgeColor',[1 0 0]); %
        rectangle('Position',[0.04+m,ymax,1,thickness],'FaceColor','w','EdgeColor',[1 0 0]); % 11 periode plafond
        rectangle('Position',[0.05+m,ymax,1,thickness],'FaceColor','r','EdgeColor',[1 0 0]); %
        rectangle('Position',[0.06+m,ymax,1,thickness],'FaceColor','w','EdgeColor',[1 0 0]); %
        rectangle('Position',[0.07+m,ymax,3,thickness],'FaceColor','r','EdgeColor',[1 0 0]); %
end
    
% 110 x 2 = 220 distance max


for p = 1:10
        m = (p-0.01)*0.1-0.1-0.35; % -0.35 translation pour faire le zéro
        rectangle('Position',[m,-thickness,3,thickness],'FaceColor','w','EdgeColor',[1 0 0]);   %
        rectangle('Position',[0.03+m,-thickness,1,thickness],'FaceColor','r','EdgeColor',[1 0 0]); %
        rectangle('Position',[0.04+m,-thickness,1,thickness],'FaceColor','w','EdgeColor',[1 0 0]); % 11 periode plafond
        rectangle('Position',[0.05+m,-thickness,1,thickness],'FaceColor','r','EdgeColor',[1 0 0]); %
        rectangle('Position',[0.06+m,-thickness,1,thickness],'FaceColor','w','EdgeColor',[1 0 0]); %
        rectangle('Position',[0.07+m,-thickness,3,thickness],'FaceColor','r','EdgeColor',[1 0 0]); %
        rectangle('Position',[0.10+m,-thickness,3,thickness],'FaceColor','w','EdgeColor',[1 0 0]);
end
        rectangle('Position',[0.625,-thickness,3,thickness],'FaceColor','w','EdgeColor','w');


GrayWidth = 0.8; % largeur du motif du miroir
%Couleur_M = [0.5 0.5 0.5]; % gris
Couleur_M = [0 0 1]; % bleu

for p = 0 : 22
       m = 0.625/11*p+0.625;
       plot([m m+0.625/11],[0 -thickness],'-','LineWidth',GrayWidth,'Color',Couleur_M)
end
plot([0.625 1.65], [0 0],'-','LineWidth',GrayWidth,'Color',Couleur_M)
plot([0.625 1.65], [-thickness -thickness],'-','LineWidth',GrayWidth,'Color',Couleur_M)
plot([0.625 0.625], [-thickness 0],'-','LineWidth',GrayWidth,'Color',Couleur_M)
plot([1.3 1.3], [-thickness 0],'-','LineWidth',GrayWidth,'Color',Couleur_M)

% for p = 7 : 20
%         m = (p-0.01)*0.1-0.1;
%         rectangle('Position',[m,-thickness-ymax,3,thickness],'FaceColor','w','EdgeColor',RGB);   %
%         rectangle('Position',[0.03+m,-thickness-ymax,1,thickness],'FaceColor',RGB,'EdgeColor',RGB); %
%         rectangle('Position',[0.04+m,-thickness-ymax,1,thickness],'FaceColor','w','EdgeColor',RGB); % 11 periode plafond
%         rectangle('Position',[0.05+m,-thickness-ymax,1,thickness],'FaceColor',RGB,'EdgeColor',RGB); %
%         rectangle('Position',[0.06+m,-thickness-ymax,1,thickness],'FaceColor','w','EdgeColor',RGB); %
%         rectangle('Position',[0.07+m,-thickness-ymax,3,thickness],'FaceColor',RGB,'EdgeColor',RGB); %
% end

couleur = [1 0 0 0.1];

for p = 1:10
        m = (p-0.01)*0.1-0.1+0.625; % +0.35 translation pour faire le zéro
        rectangle('Position',[m,-thickness-ymax,3,thickness],'FaceColor',couleur,'EdgeColor',couleur);
        rectangle('Position',[0.03+m,-thickness-ymax,1,thickness],'FaceColor','w','EdgeColor',couleur);
        rectangle('Position',[0.06+m,-thickness-ymax,1,thickness],'FaceColor',couleur,'EdgeColor',couleur);
        rectangle('Position',[0.07+m,-thickness-ymax,1,thickness],'FaceColor','w','EdgeColor',couleur);
        rectangle('Position',[0.08+m,-thickness-ymax,1,thickness],'FaceColor',couleur,'EdgeColor',couleur);
        rectangle('Position',[0.09+m,-thickness-ymax,3,thickness],'FaceColor','w','EdgeColor',couleur);
        rectangle('Position',[0.10+m,-thickness-ymax,3,thickness],'FaceColor',couleur,'EdgeColor',couleur);
end
        
        %plot([0.58 0.625],[-thickness-ymax -thickness-ymax],'k','LineWidth',0.5)
        %plot([0.58 0.625],[-ymax -ymax],'w','LineWidth',0.6)
        %plot([0.599 0.599],[-thickness-ymax+0.0015 -ymax],'w','LineWidth',0.6)
box on
        % Effacement de 2 traits noirs de la box
        plot([-0.35 -0.35],[-thickness -ymax],'w','LineWidth',0.6)
        plot([-0.35 0.625],[-0.02-ymax -0.02-ymax],'w','LineWidth',0.6)
        plot([0.625 1.65],[-0.02-ymax -0.02-ymax],'Color',couleur,'LineWidth',0.6)
        
        % traits verticaux gris
        plot([1.65 1.65],[-0.02-ymax 0],'Color',[0.5 0.5 0.5],'LineStyle','-','LineWidth',0.5)
        %plot([0 0],[0.02 ymax*0.75],'k','LineStyle','--')
        plot([0.625 0.625],[-0.02-ymax 0],'Color',[0.5 0.5 0.5],'LineStyle','-','LineWidth',0.5)
     
shadedErrorBar(x_shade,DMB_MBZ',{@nanmedian,@madX},{'-ok','markerfacecolor','k','MarkerSize',4});
axis([-0.35 1.65 -0.02-ymax ymax+0.02])
%xticks([0 0.625 1.3]) % positions des ticks sur l'abscisse
xticks([ ])
yticks([0 0.09 0.14 ymax])% positions des ticks sur l'ordonné
yticklabels({'0','0.09','0.14','0.71'}) % noms des ticks
title('1st flight recorded in half mirror on the floor condition','FontWeight','normal')%titre du graph
%xlabel('Distance travelled (m)')
ylabel('Altitude (m)')
str='     n=14';
text(0.04-0.35,ymax-0.25,str) %-0.35 translation pour faire le zéro
text(0.005-0.35,ymax-0.10,'(Bii) Impoverished visual condition') %-0.35 translation pour faire le zéro
axe = gca;
axe.FontSize=5;


% Passage à l'impression en .png de la figure
print -f9 -dpng -r1200 Demi_Miroir_Bas
% -f9 numéro de la figure
% -dpng format de la figure
% -r600 résolution
% ensuite c'est le nom

return %point d'arrêt

%faire le test de normalité
%H0 : les valeurs sont normalement distribuées
[h,p,adstat,cv] = adtest(DMB_MBZ(1,:)) %h=1 on rejette l'hypo nulle 
[h,p] = kstest(DMB_MBZ(1,:))

%faire le test de normalité de Shapiro-Wilk
for i=1:14
[H(i), pValue(i), W(i)] = swtest(DMB_CCZ(:,i))
end


%test d'égalité des var 
[h,p,ci,stats] = vartest2(DMB_MBZ(1,:),DMB_CCZ(1,:)) %h=0 cad meme var et distr normale
[h,p,ci,stats] = vartest2(DMB_MBZ(12,:),DMB_CCZ(12,:))
%test de student pour comparer chaque moyenne Two-Sample t-Test for Equal Means
[h,p,ci,stats]=ttest2(DMB_MBZ(1,:),DMB_CCZ(1,:))
[h,p,ci,stats]=ttest2(DMB_MBZ(13,:),DMB_CCZ(13,:))

%Test t pour des moyennes égales sans supposer des variances sont égales 
%t-Test for Equal Means Without Assuming Equal Variances

[h,p] = ttest2(DMB_MBZ(1,:),DMB_CCZ(1,:),'Vartype','unequal') % h=0 on ne rejette pas l'hypothèse nulle au risque 5%
[h,p] = ttest2(DMB_MBZ(11,:),DMB_CCZ(11,:),'Vartype','unequal')
```
